# Breastmilk antibody isotypes differentially protect against neonatal rotavirus infection and modulate the timing and clonal architecture of offspring B cell responses

**DOI:** 10.1101/2024.09.20.614185

**Authors:** Konjit Getachew Muleta, Simone Isling Pærregaard, Sharné Van Dijl, Giorgia Montano, Dasiel Castellanos Pérez, Rachael FitzPatrick, Asma Sadik Mustafa, Niklas Segrén, Manar Alyamani, Gina Lindberg, Gabriel Núñez, Joan Yuan, Katharina Lahl

**Author notes:** Correspondence: Katharina Lahl and Joan Yuan. equal contribution (Konjit Getachew Muleta, Simone Isling Pærregaard).

## Abstract

Mother-to-offspring transfer of antibodies protects newborns during the critical phase of early life, preventing both infection and inflammation. Whether and how pre-existing passive immunity actively shapes immune memory formation in neonates remains unclear. Using a murine neonatal Rotavirus infection model in combination with cross-foster approaches and genetic models, we disentangled the contribution of maternal IgA and IgG to neonatal immune outcomes. While breastmilk antigen specific IgA is responsible for protection from Rotavirus infection, antigen specific IgG delays pup-intrinsic humoral immune induction to after weaning. Importantly, pre-existing maternal immunity delays and constrains, rather than ablates, long term humoral immunity. Together, our data move beyond a binary view of passive immunity as protective versus inhibitory, providing a more nuanced understanding in which breastmilk antibodies actively program the neonatal response for optimal immediate protection and long-term immune education in the growing organism.

## INTRODUCTION

The event of birth marks a switch from life under sterile conditions to that under massive microbial and environmental pressure.^1,2^ The neonatal immune system is naïve to environmental challenges and newborns rely on the transfer of immunity from their mothers, which is achieved through antibodies transferred via the placenta and, after birth, through breastmilk.^3–5^ Secretory antibodies in breastmilk play important roles in establishing intestinal homeostasis through interacting with luminal microbiota and protecting from infection through immune exclusion mechanisms.^6–9^ These effects are primarily provided by secretory IgA, which is highly abundant in breastmilk, but protection from infection can also occur via breastmilk IgG, as recently shown using a mouse model of *Citrobacter rodentium* infection.^5^

The decline of maternal antibodies coinciding with solid food intake triggers an exponential increase in microbiota diversity and load, creating a “window of opportunity” during which the integration of environmental triggers by the neonatal immune system is critical for healthy imprinting of long-lasting immune homeostasis.^2,10,11^ We have shown previously that a large fraction of the IgA^+^ plasma cells (PCs) found in the adult small intestinal lamina propria (SILP) in mice is derived from B cells of early-life origin, suggesting that the PC pool in the gut retains local memory to immune triggers exposed to during early life.^12^ Breastmilk antibodies, particularly of the IgG isotype, can actively suppress humoral immune activation to the microbiota during the time of weaning, and withdrawal was shown to predispose mice to inflammatory bowel disease onset caused by a microbiota-directed immune overreaction.^8,9^ Breastmilk IgA can contribute to the delay of anti-microbial pup-intrinsic IgA induction, presumably through its role in immune exclusion and the ensuing blockade of microbial dissemination to immune inductive sites.^4^ Under specific pathogen free conditions, pups start producing secretory IgA at the time of weaning, correlating with the cessation of breastmilk antibody exposure and the maturation of gut-associated lymphoid tissues.^4^

In contrast to steady-state humoral immune induction at the intestinal wall, Rotavirus (RV) infection in pups born to RV-naïve dams induces a strong, long-lasting IgA response already during the first week of life.^13^ Previous studies from human vaccine trials reported that RV-specific IgA in breastmilk negatively correlated with seroconversion in the babies shortly after immunization, suggesting that breastmilk antibodies might be inhibitory of pup-intrinsic immunity also in response to viral triggers.^14–16^ The effects however were small and, given the endemic nature of the virus and the high levels of immunity in the human population, less consistent than one would predict.

To gain a better idea of the effects that pre-existing maternal immunity might have on humoral immune activation in the offspring in the context of a potent viral infection, we here analyzed humoral immune onset and longevity in pups raised by RV-immune dams and exposed to murine RV as neonates. We found that humoral immune memory induction occurs in offspring despite protection from infection by RV-specific maternally derived IgA, leading to efficient protection from reinfection during adulthood. In contrast to pups nursed by RV-naïve dams, however, RV-specific humoral immunity is delayed until after weaning, and the delay was mediated by maternal RV-specific IgG. Our findings challenge the dogma that the active role in immune priming by maternal antibodies is restricted to immune system inhibition in suckling offspring and have important implications for understanding general early-life immune imprinting mechanisms and rational vaccine design.

## RESULTS

### Maternal immunity protects suckling pups from Rotavirus infection without inhibiting functional immune memory establishment

The murine Rotavirus (RV) strain EC_W_ causes acute asymptomatic infection in adult mice and leads to self-limiting diarrhea in pups. Regardless of age, RV is cleared 7-10 days post infection in wildtype (WT) mice.^17–19^ To quantify the impact that maternal immunity has on protection and immune induction in the suckling offspring, we orally infected pups born and raised by immune (infected with RV three weeks prior to conception) or naïve dams with RV (Fig. 1A). We detected few viral particles in a subset of pups born to immune dams at day 3 and no virus could be detected in the intestinal content from these pups at day 5 post exposure (Fig. 1B). In addition, no pups from immune dams developed diarrhea (0/27 versus 12/16 pups from naïve dams). This confirms that mother-derived immune mechanisms efficiently curb the viral load in the suckling offspring.

**Figure 1:**
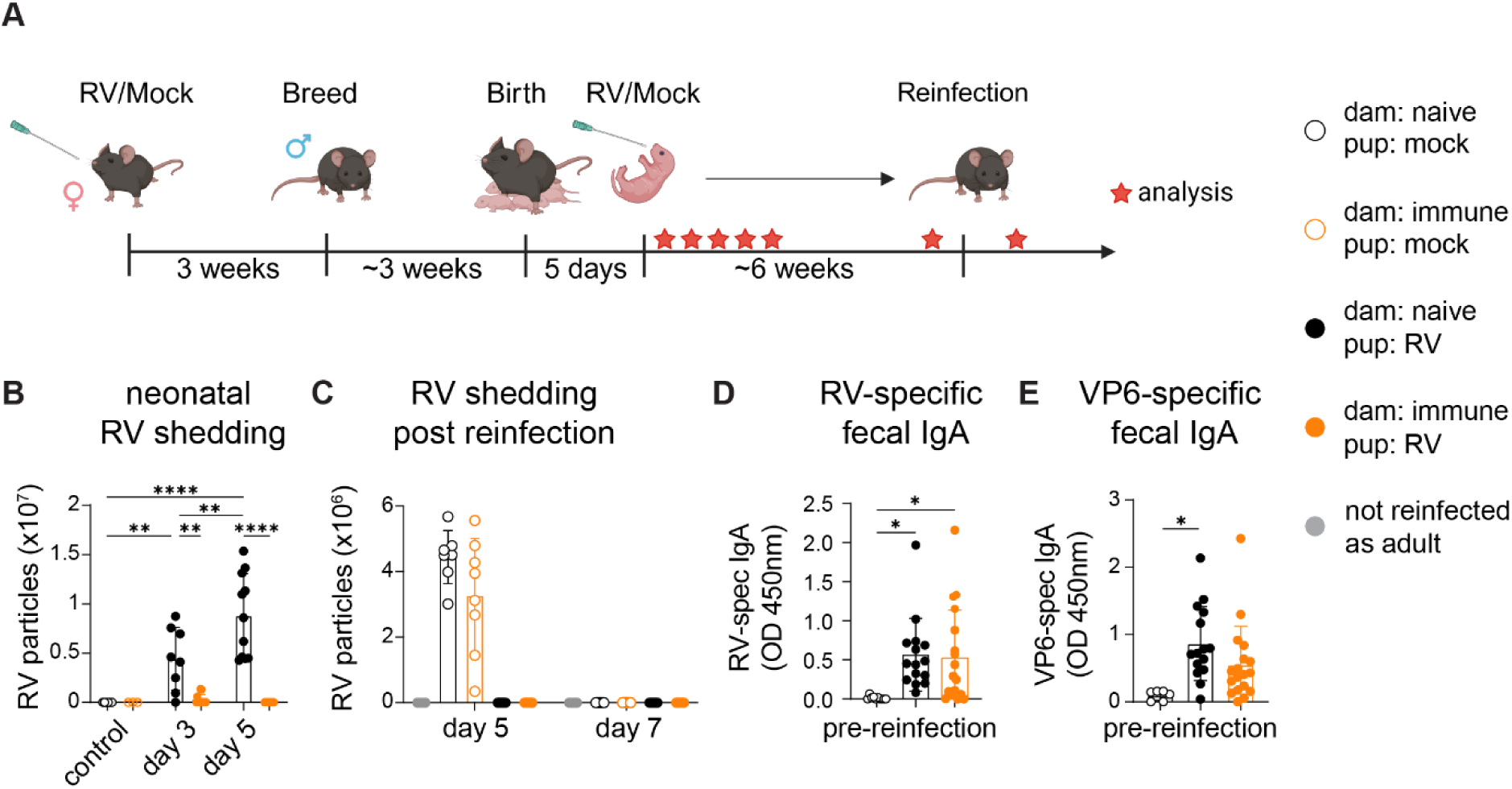
Maternal immunity protects suckling pups from Rotavirus infection without inhibiting immune memory induction. A) Experimental outline. B) RV shedding in the colon content of orally RV infected pups at five days of age born to either naïve (black) or immune (orange) dams. ELISAs were performed at three and five-days post infection. Data from 3 independent experiments with one litter per group (n = 2-7/repeat). Repeats are from separate litters per experimental condition, distributed over the two analysis time points. C) RV shedding in feces collected from mice orally infected with RV at five days old and re-infected at 6 weeks post primary infection. Fecal samples were collected at five and seven days post reinfection for ELISAs. The graph represents two experiments with n = 2-5 mice/repeat. D) RV-specific fecal IgA ELISA of samples collected at 5-8 weeks post infection from mice orally infected with RV at five days old. The graph shows three experimental repeats with n = 2-7 mice/repeat. E) VP6-specific IgA ELISA on fecal samples collected at 5-8 weeks post infection from mice orally infected with RV at five days old. The graph shows three independent experiments (n = 2-7/repeat). Each data point represents an individual mouse, bars depict mean +/- SD. Statistical analyses were performed using ordinary one-way ANOVA with Tukeýs multiple comparison test. *p < 0.05, **p < 0.01, ***p < 0.001, ****p < 0.0001.

We reasoned that highly effective protection may interfere with immune induction in the pup at an early age, in line with findings in human studies showing that RV-specific breastmilk IgA correlated with poor seroconversion in babies upon oral vaccination with the attenuated live RV vaccine Rotarix^14–16^, which has recently been confirmed in a mouse model.^20^ However, offspring protected from productive infection upon oral RV exposure by maternal immune factors were equally protected from reinfection post weaning as those mice that showed symptomatic infection and measurable RV titers as pups, demonstrating that maternal immunity does not preclude the establishment of functional immunological memory to the WT EC_W_ RV strain (Fig. 1C). Accordingly, despite the vastly different levels of viral shedding following infection, amounts of fecal RV-specific IgA did not significantly differ in adult mice regardless of whether they were productively infected as neonates or not (Fig. 1D). Likewise, these mice had similar amounts of IgA against the immunodominant epitope, virus protein (VP)6, despite this epitope being part of the middle capsid layer and only exposed after cellular entry (Fig. 1E). We conclude that maternal immune factors confer protection of suckling offspring without prohibiting functional immune memory establishment capable of protecting from subsequent post weaning viral challenge.

### RV-exposure in maternally protected pups induces RV-specific humoral immunity with delayed kinetics

We next interrogated whether maternal protection impacted the kinetics of the early-life induced immune response to RV, given its dramatic effect on viral load. Interestingly, maternally protected pups showed mesenteric lymph node (mLN) hypertrophy seven days post exposure to RV, a general sign of immune exposure (Fig. 2A).^21^ This included an overall increase in B cells, 14 days post infection (Fig. 2B, see Suppl. Fig. 1A for gating strategy). Overall isotype switched B cells and plasmablasts were highly variable in the mLN 14 days post infection (Fig. 2B), and IgA^+^ plasma cells (PCs) in the small intestinal lamina propria (SILP) of maternally protected pups were only marginally increased over those in uninfected controls, while the intestines of RV-infected pups from naïve mothers harbored significantly more IgA^+^ PCs (Fig. 2C, see Suppl. Fig. 1B for gating strategy). Lastly, we looked for the occurrence of RV-specific IgA^+^ PCs using virus-like particles (VLP) constructed of the VP6 protein fused to VP2-GFP. Surprisingly, we were unable to detect any RV-specific IgA^+^ plasmablasts in the mLN and PCs in the SILP 14 days post neonatal infection in the presence of maternal protection (Fig. 2D), despite the abundance of anti-VP6 IgA in adulthood (Suppl. Fig. 1C).

**Figure 2:**
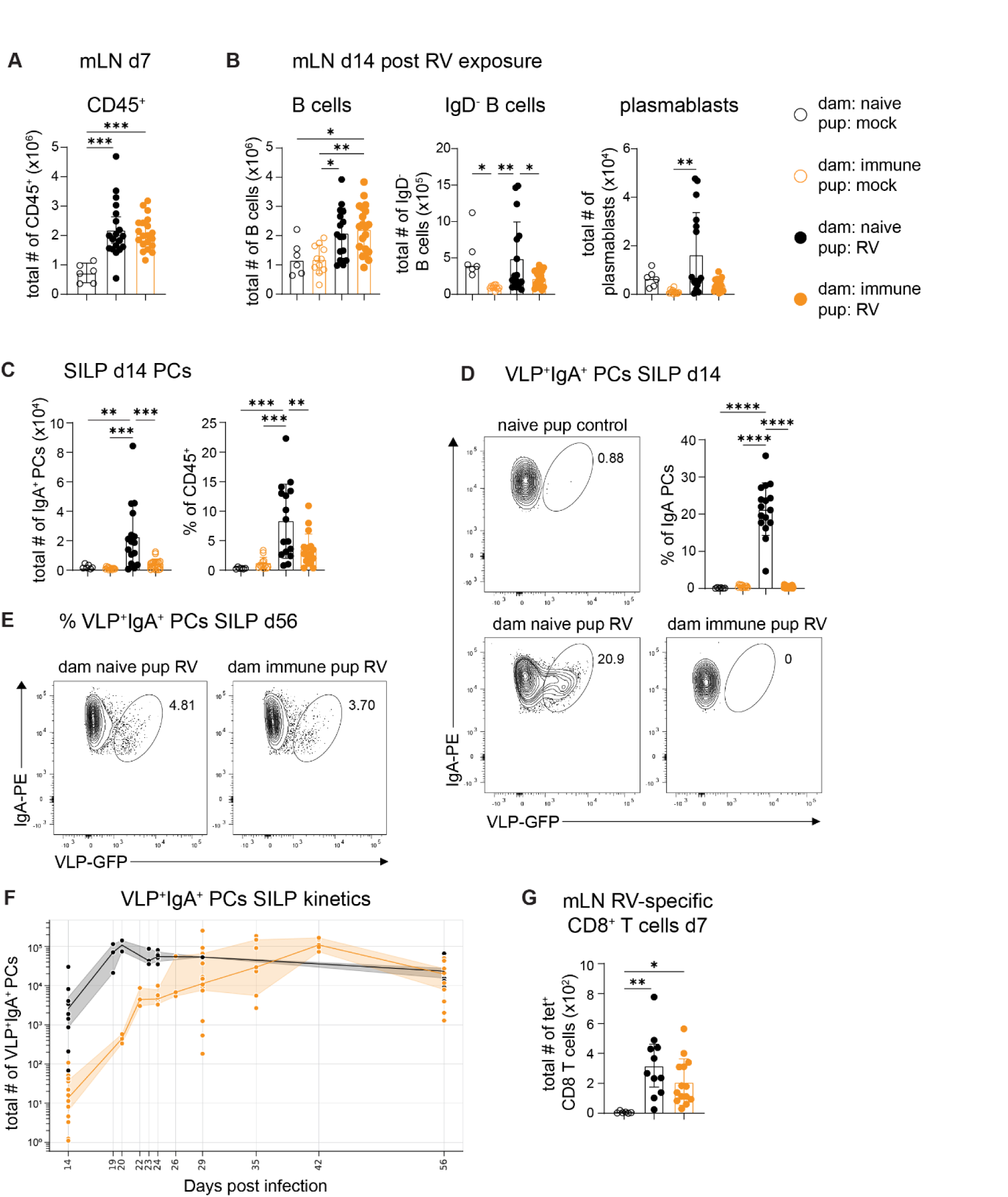
RV-exposure in maternally protected pups induces RV-specific humoral immunity with delayed kinetics. A) Mesenteric lymph node (mLN) cellularity (measured as single cell, live, CD45^+^ cells) 7 days post neonatal day 5 oral RV infection. The graph shows 4 independent experiments with n = 3-8 mice per litter except for the naïve groups where data is from 2 independent experiments, n = 3/repeat. B) Total numbers of mLN B cells (single cells, live CD45^+^ CD19^+^), mature B cells (single cells, live CD45^+^CD19^+^IgD^-^) and plasmablast (single cells, live CD45^+^CD19^+^B220^-/low^IgD^-^CD138) analyzed two weeks post neonatal day 5 oral RV infection. Data is from 3 or 4 experiments with n = 1-8/repeat, except the naïve dam, naïve pup group, which is from one litter with 6 pups. C) Total numbers (left) and frequencies (right) of small intestinal lamina propria IgA^+^ plasma cells (PCs) (single cells, live CD45^+^CD98^+^B220^low/-^IgA^+^) analyzed two weeks post neonatal day 5 oral RV infection. 3-4 experiments with n = 1-8/repeat, except the naïve dam, naïve pup group, which is from one litter with 6 pups. D) Representative FACS plots and frequency of RV-specific (VLP^+^) IgA^+^ PCs in the SILP analyzed two weeks post neonatal day 5 oral RV infection. The graph shows VLP^+^ IgA^+^ PCs as a percentage of IgA^+^ PCs from 3-4 experiments with n = 1-7/repeat, expect the naïve dam, naïve pup group, which is from one litter with 6 pups. E) Representative FACS plots showing RV-specific (VLP^+^) IgA^+^ PCs at 8 weeks post neonatal day 5 oral RV infection. F) Absolute cell counts of VLP^+^IgA^+^ SILP PCs at indicated time points post infection. All mice were orally infected 5 days after birth. Naïve dam: 14 dpi (3 experiments); 19, 20, 23, and 24 dpi (1 experiment each); 56 dpi (2 experiments). Immune dam: 14 dpi (5 experiments); 20, 22, 24 and 26 dpi (1 experiment each); 29 dpi (4 experiments); 36 dpi (2 experiments); 42 dpi (1 experiment); 56 dpi (3 experiments). G) Absolute cell numbers of RV specific (VP6 tetramer^+^) CD8^+^ T cells in the mLNs of pups 7 days post neonatal day 5 oral RV infection. The graph shows three independent experiments with n= 3-6/repeat, except for the naïve controls, which were performed twice with n= 3/repeat. Each data point represents an individual mouse. Open circles represent uninfected pups, full circles represent infected pups. Black circles represent pups from naïve dams and orange circles represent pups from immune dams. (A-G) Bars depict the mean +/- SD. Statistical analyses were performed using ordinary one-way ANOVA with Tukeýs multiple comparison test. *p < 0.05, **p < 0.01, ***p < 0.001, ****p < 0.0001. (F) Line depicts the median and shades represent the 95% confidence interval.

To better understand the discrepancy between the pre- and post-weaning anti-RV response, we performed an in-depth kinetic analysis of RV-specific humoral immune onset in the SILP. We found that VP6-specific humoral immunity is robustly detectable 14 days post infection in offspring of naïve dams, peaks around three weeks post infection, and remains relatively stable for at least seven weeks post infection (Fig. 2E, F). In contrast, RV-specific VLP^+^IgA^+^ PCs in pups protected by pre-existing maternal immunity were scarce before weaning, but rose around three to four weeks post infection, showing some variability in the timing of onset, but largely converging in abundance with those found in pups from naïve dams around four to five weeks post infection (Fig. 2E, F).

Interestingly, the detected blockade of pre-weaning antigen-specific immune onset by pre-existing maternal immunity was specific to the B cell response, as RV-specific CD8 T cells developed comparably in maternally protected and unprotected pups (Fig. 2G, see Suppl. Fig. 1D for gating strategy), ruling out a general deficit of RV-specific adaptive immune induction in maternally protected pups.

### Pre-existing maternal immunity delays RV-specific humoral immunity without impairing overall germinal center (GC) formation

The bulk of the RV-specific IgA response is T cell dependent and arises from a GC response.^22^ To determine whether maternal immunity acts by blocking GC formation or by impairing downstream PC differentiation, we compared the GC reaction in RV-exposed pups from naïve and immune dams 14 days post infection by quantifying overall T follicular helper cells (Tfh) and GC B cells in the mLN. Maternally protected pups mounted a robust overall GC response upon neonatal RV exposure (Fig. 3A), consistent with the observed mLN hypertrophy (Fig. 2A, B). However, RV-specific GC and Tfh responses were severely blunted in maternally protected pups at this time point (Fig. 3B, gating strategy Suppl. Fig. 1A and 2A, respectively). These results demonstrate that pre-existing maternal immunity acts selectively on RV-specific B cell responses, suppressing antigen-driven GC seeding or expansion, without broadly suppressing GC formation.

**Figure 3:**
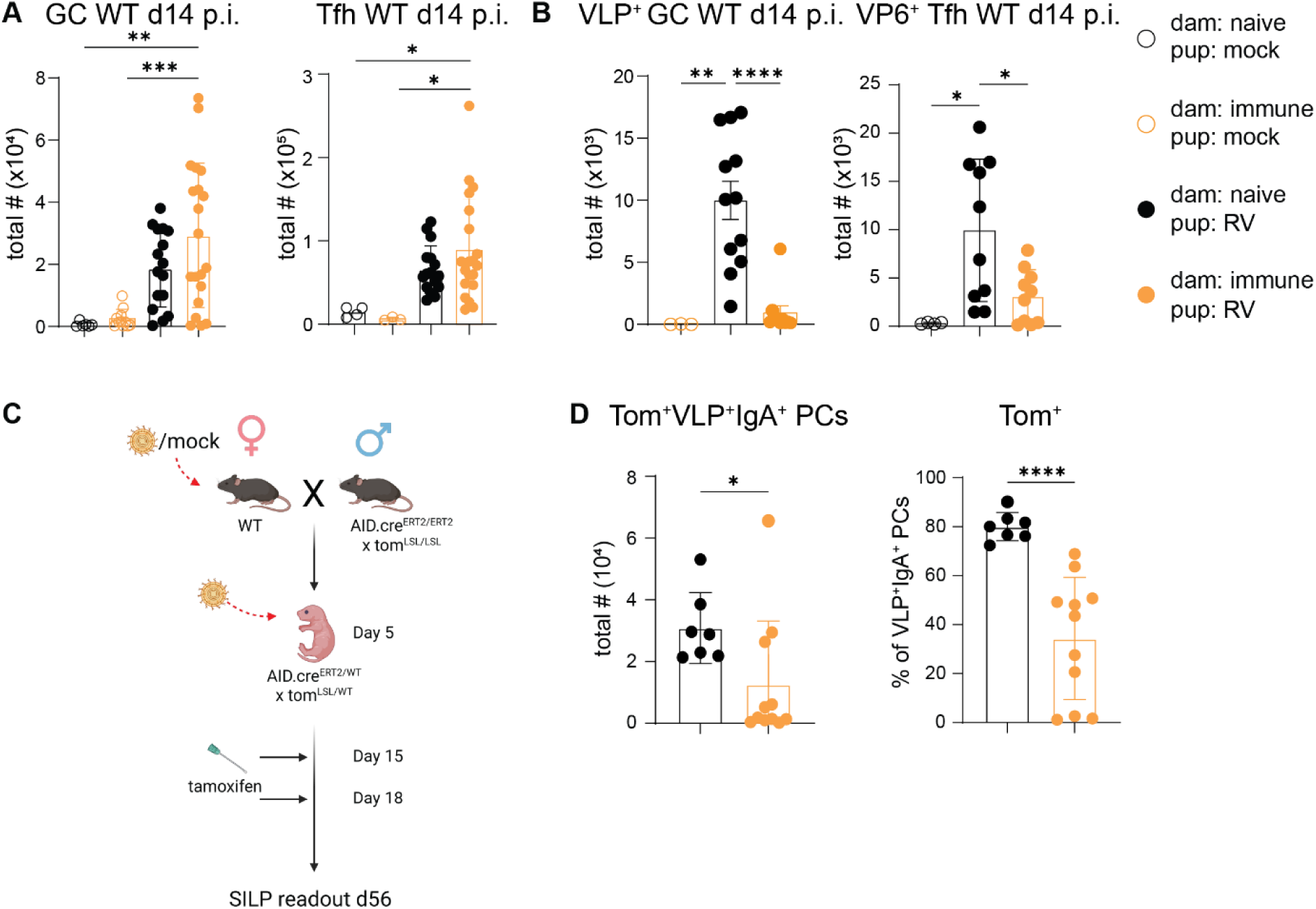
Temporary blockade of humoral immunity in the context of maternal protection occurs at the germinal center (GC) level. A) Total Germinal Center (GC) B-cell (left) (single cells, live CD3^-^Gr1^-^B220^+/-^IgD^-^CD19^+^CD95^+^GL7^+^) numbers and total T follicular helper cell (right) (single cells, live CD3e^+^B220^-^CD4^+^CD44^+^PD-1^+^Bcl6^+^) numbers 2 weeks post neonatal day 5 RV infection of pups born to naïve (full black circles) or immune (full orange circles) dams, non-infected pups (orange/black open circles if born to immune/naive dams, respectively). The GC B-cell graph shows 1 experiment, n=6, for dam naive – pup naïve; 3 experiments, n=3-4/repeat, for dam immune – pup naïve; 4 experiments, n=1-6/repeat for dam naïve – pup RV; and 4 experiments for dam immune – pup RV, n = 4-8/repeat. The Tfh cell graph shows 1 experiment, n=4, for dam naive – pup naïve; 1 experiment, n= 3, for dam immune – pup naïve; 4 experiments, n=3-6/repeat for dam naïve – pup RV; and 5 experiments for dam immune – pup RV, n = 3-6/repeat. B) Total VLP^+^ GC B-cell numbers (left) and total RV-specific (VP6 tetramer^+^) Tfh cell numbers (right) 2 weeks post postnatal day 5 RV infection of pups born to naïve (full black circles) or immune (full orange circles) dams, or naïve pups (orange/black open circles if born to immune/naive dams, respectively). Left graph shows 3 experiments for each RV infected group and one experiment for the dam immune - pup naïve control (n=3-6/repeat). Right graph shows 2-3 experiments (dam naïve – pup RV: 3 experiments, n=3-4/repeat; dam immune – pup RV: 2 experiments n=4-6/repeat; dam naïve - pup naïve: 1 experiment, n= 4). C) Experimental outline of time-stamping experiment. D) Total Tomato^+^VLP^+^IgA^+^ SILP PC numbers (left) and frequencies of Tomato^+^ as a percentage of VLP^+^IgA^+^ SILP PCs (right) 5-8 weeks post neonatal day 5 RV infection. The graph shows pooled data from 4 experiments with 1-4 mice per repeat (all repeat on individual litters). Each data point represents an individual mouse. (A-B, D) Bars depict mean +/- SD. Statistical analyses were performed using (A-B) ordinary one-way ANOVA with Tukeýs multiple comparison test, (D) Two-tailed Mann-Whitney test. *p < 0.05, **p < 0.01, ***p < 0.001, ****p < 0.0001.

To interrogate the origin of RV-specific PCs in maternally protected pups, we employed *Aicda*-Cre^ERT2^ inducible fate-mapping to permanently label B cells with a history of GC participation at the time of tamoxifen administration. Pups were generated by crossing immune or naïve WT dams with double homozygous *Aicda*-Cre^ERT2^ Rosa26^TdTomatoLSL^ sires and RV-infected on day 5 of postnatal life. Tamoxifen was administered at 10 and 13 days post infection to fate-map activated B cells upon robust GC formation (Fig. 3C). Near complete labeling of total GC B cells was observed in both groups at two weeks post infection (Suppl. Fig. 2B), confirming efficient labelling. However, when we examined the long-term antigen specific GC output by quantifying the extent of tomato labeling among SILP VLP^+^IgA^+^ PCs at 5-8 weeks post infection, we observed a significant difference. While 80% of adult RV-specific IgA^+^ PCs in offspring from naïve dams were labeled, maternally protected offspring showed reduced and much more variable labelling frequency, demonstrating that fewer antigen-specific B cells had successfully become activated or expanded within the GC during the labelling window (Fig. 3D, for gating strategy see Suppl. Fig. 2C). Together, these data demonstrate that the delay in RV-specific humoral immune induction occurs upstream of PC differentiation, at the level of antigen-specific B cell priming or GC expansion.

### RV-specific IgA^+^ PCs from maternally protected offspring show reduced clonal diversity

To assess whether clonal diversity of RV-specific IgA^+^ PCs differed between maternally protected and unprotected offspring, we sorted SILP VLP⁺IgA⁺ PCs from 5-8 week old neonatally infected mice born and raised by immune or naïve dams and performed bulk IgHA VDJ repertoire sequencing. The total number of SILP VLP⁺IgA⁺ PCs was comparable between the two groups (Supplementary Fig. 3A) ensuring comparable sequencing depth. Offspring of immune dams showed a markedly reduced clonal diversity, characterized by restricted VH-JH gene segment usage (Fig. 4A) and the dominance of a limited number of clones (Fig. 4B). This is reflected both in reduced total clone counts (Fig. 4C) and a lower Shannon diversity index (Fig. 4D and Suppl. Fig 3B). These data demonstrate that pre-existing maternal immunity significantly constrained the breadth of the antigen-specific IgA response against the immunodominant RV epitope. Strikingly, despite the observed reduction in early GC participation of RV-specific clones, the frequency of somatic hypermutations (SHM) per VLP^+^ clone was not significantly altered in offspring of immune dams (Fig. 4E), indicating that clones once successfully recruited into the GC undergo extensive affinity maturation. Taken together, these results demonstrate that the recruitment of antigen-specific founder B cells into the GC reaction is constrained without impairing the quality of GC function in maternally protected offspring.

**Figure 4:**
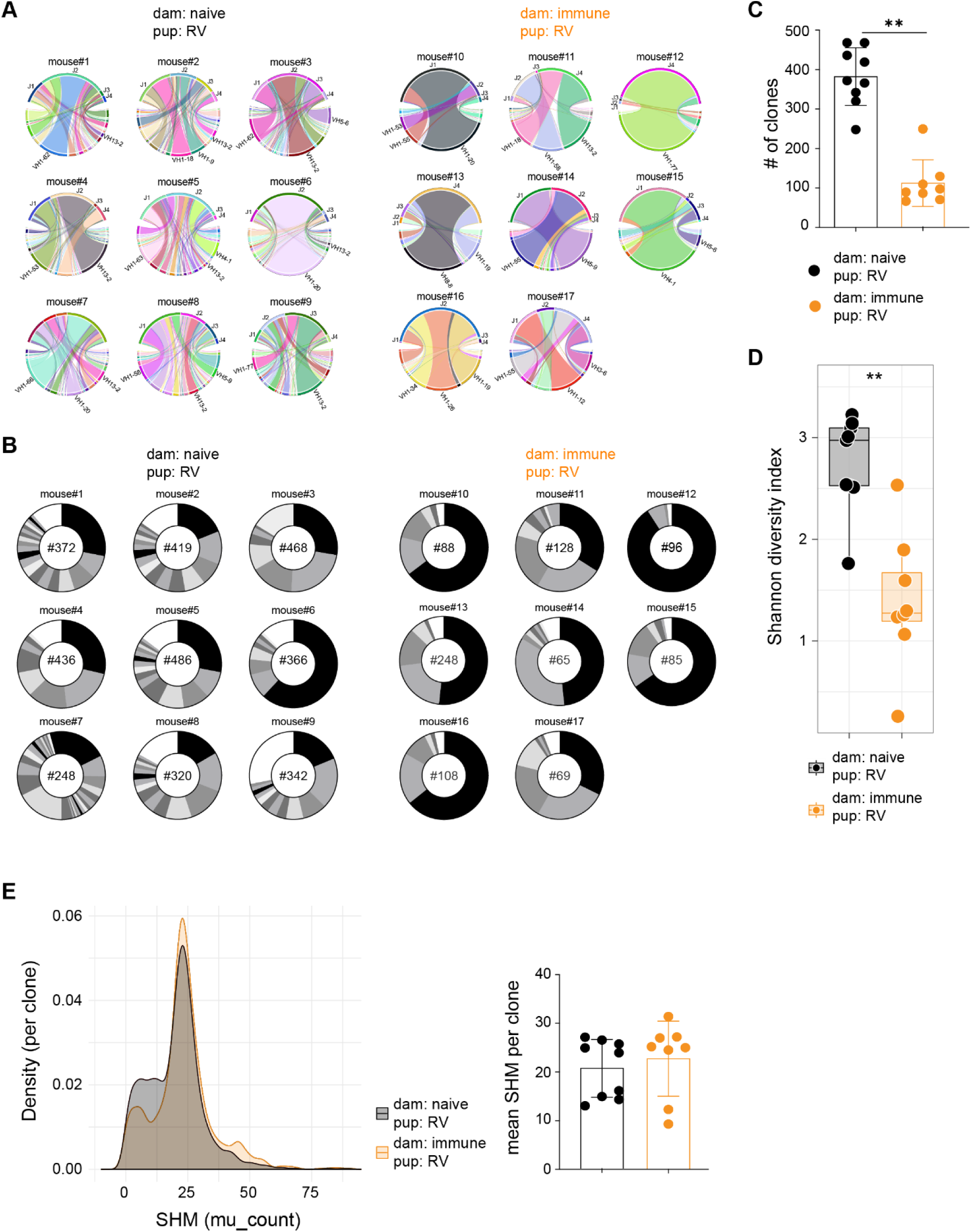
PCs from maternally protected offspring show less clonal diversity. A) VLP^+^IgA^+^ SILP PCs were FACS-sorted from adult offspring 5-8 weeks following neonatal day 5 RV infection. Sorted cells were subjected to bulk IgHA VDJ repertoire sequencing. Circos plots show VH-JH gene segment pairing frequencies from all samples born to naïve dams (left, n=9 mice) or immune dams (right, n=8 mice) following oral RV infection neonatal day 5. Each chord represents a unique VH-JH combination, with chord width proportional to the frequency of the corresponding VH and JH family usage. Outer arcs indicate overall VH and JH gene segment usage. B) Donut plots showing clonal distributions per sample in adult offspring born to naïve dams (left, n=9 mice) or immune dams (right, n=8 mice) following oral RV infection neonatal day 5. Each segment corresponds to a distinct clone, and segment size reflects the percentage of total sequencing reads assigned to that clone. Clones representing less than 1% of total reads were grouped into a single segment (shown in white). The total number of clones is indicated in the middle of the donut plots. C) Quantification of unique clones per sample in adult offspring born to naïve dams (black, n = 9) or immune dams (orange, n = 8) following oral RV infection neonatal day 5. Bars indicate mean ± SEM with individual data points overlaid. D) Shannon diversity index calculated from clone frequencies within each sample of adult offspring from naïve (black, n = 9) or immune (orange, n = 8) dams 5-8 weeks post neonatal day 5 RV infection. Each dot represents one biological replicate; box-and-whisker plots indicate median and interquartile range, with whiskers showing minimum and maximum values. E) Density plot (left) showing somatic hypermutation (SHM) in the IgHA variable region across B cell clones from adult offspring born to naïve (black, n = 9) or immune (orange, n = 8) dams 5-8 weeks following neonatal day 5 RV infection. Biological replicates were combined, and mutation count distributions are shown without weighting for clone size. Mean SHM per clone for each sample in the two groups (right). (C, E) Bars depict mean +/- SEM. (C-E) Statistical significance was assessed using a two-tailed Wilcoxon rank-sum test **p< 0.01. Data is pooled from 3 independent experiments.

### Protection from RV infection and delay of RV-specific humoral immunity in maternally protected pups are orchestrated by breastmilk antibodies

Previous studies correlated maternal antigen-specific antibodies with poor seroconversion in RV vaccinated children^14–16^ and RV-specific breastmilk IgA was shown to block intrinsic RV-specific IgA production in mice upon vaccination.^20^ Indeed, the lack of B cells in previously infected dams reversed the delayed onset phenotype of SILP RV-specific IgA^+^ PCs of maternally protected pups (Suppl. Fig. 4A, B), consistent with antibody mediated transgenerational effects. Pathogen-specific maternal antibodies are delivered to the baby via the placenta (yolk sac in mice^23^) prior to birth as well as in breastmilk post birth. To test for the differential contribution of both pathways, we cross-fostered pups born to immune or -naïve dams on the first day after birth (Fig. 5A). As expected, we found that maternal protection from RV infection at day 5 after birth is mediated entirely by breastmilk antibodies from immune dams, as pups born to immune dams but raised by naïve dams showed equally high viral load compared to those born and raised by naïve dams (Fig. 5B). Importantly, the delay in RV-specific IgA^+^ mLN plasmablast and SILP PC occurrence was entirely dependent on the immune status of the foster mother, but not on that of the birth mother, as we detected negligible RV-specific IgA^+^ mLN plasmablast and SILP PC responses 14 days post-infection in pups born and raised by immune dams or born to naive dams and cross-fostered to immune dams (Fig. 5C). These data exclude a role for antibodies delivered *in utero* in regulating humoral immune onset in this model. As expected from non-fostered groups (Fig. 2A, B), the immune status of neither the birth or foster dams affected the increase in overall cellularity and B cell numbers in the mLN upon neonatal RV exposure (Suppl. Fig. 4C). As previously noted, overall IgA^+^ PC numbers were somewhat more pronounced in unprotected pups (Suppl. Fig. 4D). Together, we conclude that breastmilk antibodies are the primary mediators of maternal influence on neonatal immune outcomes upon RV infection.

**Figure 5:**
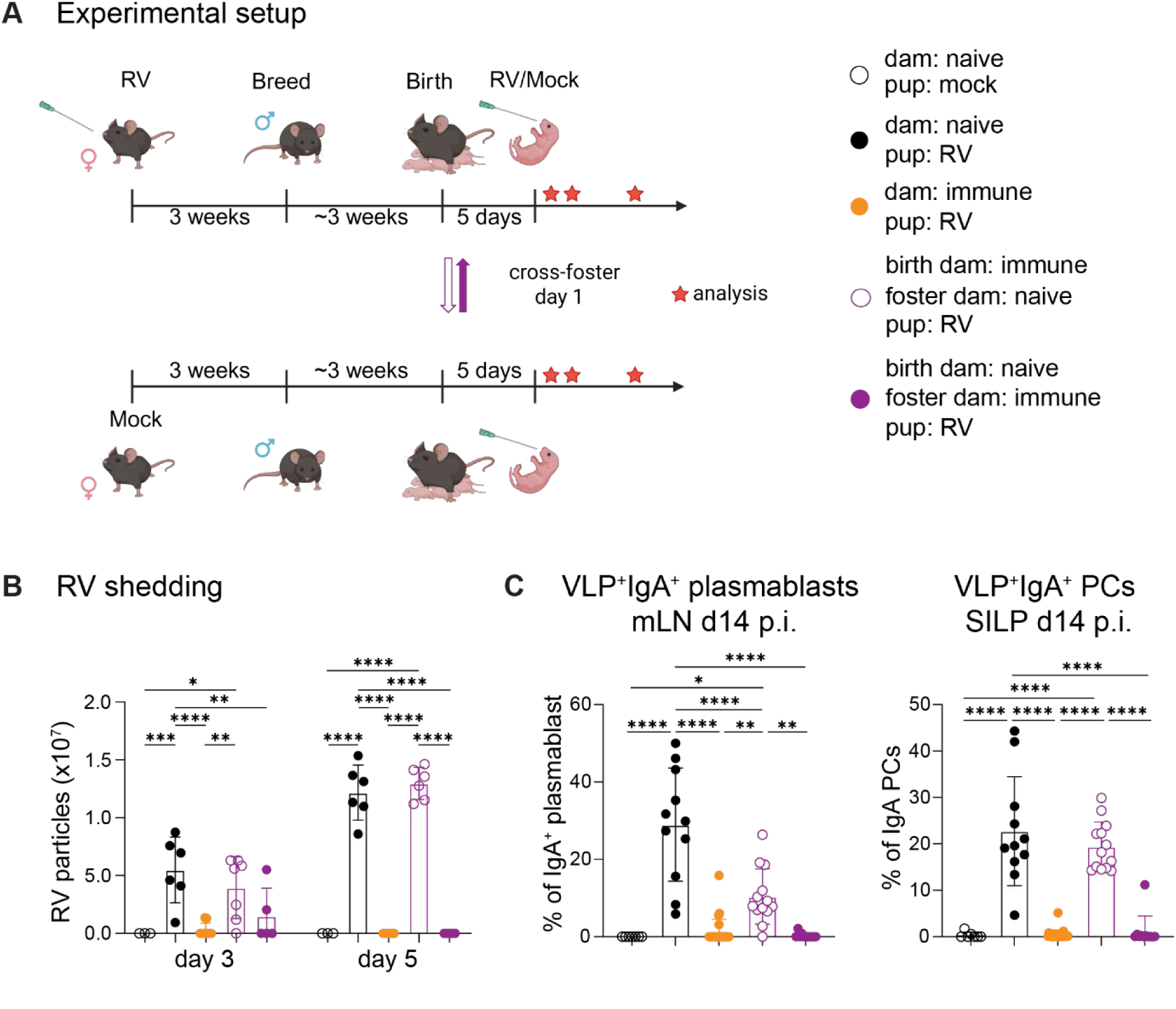
Protection from RV infection and delay of RV-specific humoral immunity in maternally protected pups are orchestrated by breastmilk antibodies. A) Experimental layout for cross foster experiments. B) RV shedding in the colon content of orally RV infected neonatal day 5 pups at three and five days post infection measured by ELISA. Cross-fostered litters were swapped within the first 24 hours post birth and were either born to immune and raised by naive (open purple circles) or born to naïve and raised by immune dams (closed purple circles). Each experimental group consists of 2 independent litters with n = 3-8 pups/litter, except for the naïve control groups that represent one experiment (n = 3). C) Frequency of RV-specific (VLP^+^) IgA^+^ plasmablasts in the mLN (left) and RV-specific (VLP^+^) IgA^+^ SILP PCs (right) analyzed two weeks post neonatal day five oral RV infection. Naive group 1 experiment n = 7, non-cross fostered groups 3-5 experiments n = 2-8/repeat, cross fostered groups 2 experiments n = 5-8/repeat. (B-C) Each data point represents an individual mouse. Bars depict mean +/- SD. Statistical analyses were performed using Ordinary one-way ANOVA with Tukeýs multiple comparison test. *p < 0.05, **p < 0.01, ***p < 0.001, ****p < 0.0001.

### Maternal RV-IgA protects pups from infection, whereas maternal RV-IgG delays the neonatal intestinal RV-specific humoral immune response

RV-specific IgA protects adult mice from reinfection through immune exclusion and intracellular neutralization^17,24,25^ and is likely to also mediate maternal protection in our model. We thus measured RV-specific IgA from the milk-plug in the stomachs of nursing pups and found that it is highly abundant in the breastmilk of immune dams (Fig. 6A). When breastfeeding RV-exposed pups, this amount increases over time, consistent with the established positive feedback loop between nursing dam and offspring^26^. Interestingly, we also detected abundant RV-specific IgG1 in the stomach content of pups born and raised by immune dams regardless of the pup’s exposure status, showing that murine breastmilk from immune dams contains ample RV-specific IgG in addition to IgA (Fig. 6B). As expected, RV-specific IgA and IgG1 in the stomach content were foster-dam derived in cross-fostered pups (Suppl. Fig. 5).

**Figure 6:**
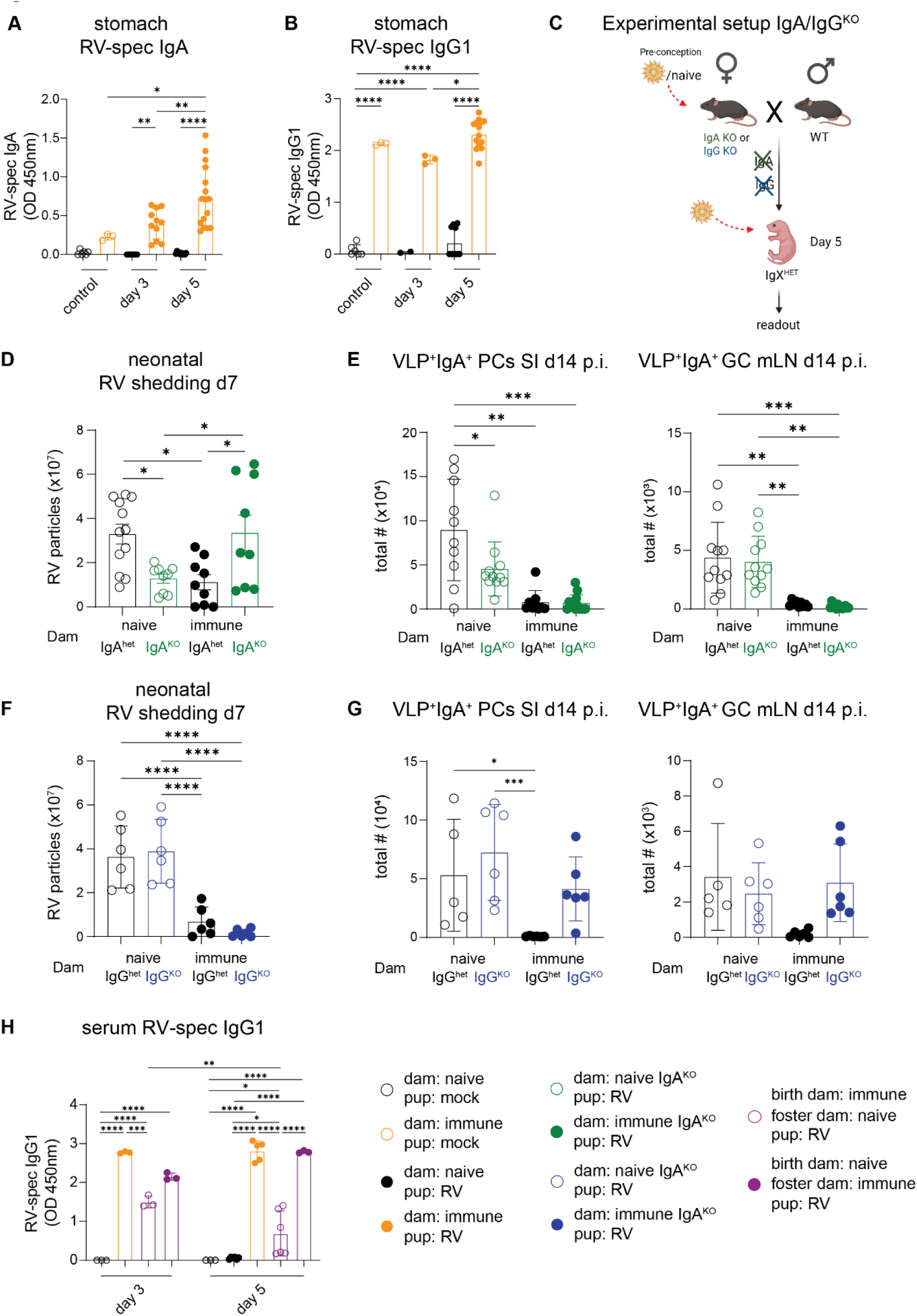
Maternal RV-IgA protects pups from infection, whereas maternal RV-IgG delays the neonatal intestinal RV-specific humoral immune response. A) RV-specific IgA in the stomach contents of RV infected (full circles) or naive (open circles) pups at neonatal day 5 born to immune (orange) or naive (black) dams three and five days post neonatal day five oral RV infection. The graph shows 2-4 independent experiments with n = 3-5 pups per repeat, except for the dam immune - pup naive (day 3) group, which was performed once with n = 3 pups. B) RV-specific IgG1 in the stomach contents of RV (full circles) or mock (open circles) infected pups born to immune or mock infected dams three and five days post neonatal day five oral RV infection. The graph shows 2-4 independent experiments with n = 3-5 pups per group, except for the following groups showing 1 experiment with n = 2-3 pups: dam naïve - pup RV (day 3, n=3), dam immune - pup RV (day 3, n=2), and dam immune – pup naïve (day 7, n=3). C) Experimental outline of experiments using IgA and IgG-deficient dams. D) RV shedding in the colon content of neonatal day 5 orally RV infected pups born to either naïve (open circles) or immune (full circles) IgA HET (black) or IgA KO (green) dams. ELISAs were performed at seven days post infection. The graph shows 2-3 independent experiments with n = 3-6/repeat, negative values have been changed to 0. E) Total numbers of VLP^+^IgA^+^ SILP PCs (left) and VLP^+^GC B-cells (right) in pups born to naïve (open circles) or immune (full circles) IgA HET (black) or IgA KO (green) dams 2 weeks post neonatal day 5 oral RV infection. The graphs show 3 experiments n = 2-3/repeat. F) RV shedding in the colon content of neonatal day 5 orally RV infected pups born to either naïve (open circles) or immune (full circles) IgG HET (black) or IgG KO (blue) dams. ELISAs were performed at seven days post infection. The graph shows 2 independent experiments with n = 3/repeat, negative values have been changed to 0. G) Total numbers of VLP^+^IgA^+^ SILP PCs (left) and VLP^+^GC B-cells (right) in pups born to naïve (open circles) or immune (full circles) IgG HET (black) or IgG KO (blue) dams 2 weeks post neonatal day 5 RV infection. The graph shows 2 independent experiments with n= 2-3/repeat. H) RV-specific IgG1 in the serum of mock (black open circles) or RV infected pups born to immune (orange) or mock infected (black) dams three and five days post neonatal day 5 oral RV infection. Cross fostered litters were swapped within the first 24 hours post birth and were either born to immune and raised by naive (open purple circles) or born to naive and raised by immune dams (full purple circles). The graph shows 2 independent experiments with (n= 2-4) for the 5 days post infection group. For the naïve control group and 3 days post infection groups, the graph shows 1 experiment with n = 3/group. Each data point represents an individual mouse. (A-G) Bars depict mean +/- SD. Statistical analyses were performed using (A, B, G) ordinary one-way ANOVA with Tukeýs multiple comparison test. (C-F) 2-way ANOVA with Tukey’s multiple comparison test. *p < 0.05, **p < 0.01, ***p < 0.001, ****p < 0.0001.

To dissect isotype specific effects of breastmilk antibodies, we obtained mice deficient in either IgA^27^ or IgG^28^. Heterozygotic (HET) or knockout (KO) littermate dams were bred with WT males ensuring pups were sufficient in IgA and IgG. We infected pups at five days of age and analyzed viral load by shedding ELISA seven days later (Fig. 6C). In line with the reported role for IgA in providing protection from reinfection with RV in adult mice^17^, we found that pups born to immune dams deficient in IgA were susceptible to RV infection (Fig. 6D). Interestingly, pups from immune IgA^KO^ and IgA^HET^ dams showed similarly blunted RV-specific IgA^+^ PC responses in the SILP and RV-specific GC B cell induction in the mLN at two weeks post infection (Fig. 6E), effectively uncoupling viral load from the temporal modulation of immune induction. In contrast, while maternal RV-IgG was dispensable for protection from neonatal RV infection (Fig. 6F), it was entirely responsible for the delay in pup intrinsic humoral immune induction, as pups from immune IgG^KO^ dams showed similar RV-specific IgA^+^ PCs in the SILP and RV-specific GC B cell induction in mLN as unprotected pups at two weeks post infection (Fig. 6G).

Antibody-mediated immune modulation is an established phenomenon^29^, but in order to actively regulate humoral response induction, breastmilk antibodies would need to cross the epithelial layer of the small intestine. Indeed, retrograde uptake of breastmilk IgG, but not IgA, has been reported before.^30,31^ To test this in our model, we measured RV-specific IgG1 in the serum of pups from our cross fosters and found that the majority was derived from breastmilk, as those pups born to naïve dams and raised by immune mothers had significantly more RV-specific IgG1 in their serum than pups born to immune mothers but raised by naïve mothers (Fig. 6H). Together, our data support a model in which RV-specific breastmilk IgA mediates luminal protection from infection, while RV-specific IgG modulates early-life humoral immune onset in the context of maternal protection.

## DISCUSSION

Here, we report that breastmilk IgA from immune dams provides efficient protection from neonatal RV infection in a mouse model. Conversely, breastmilk IgG from immune dams leads to a delay of offspring-intrinsic humoral immunity, which ultimately develops after weaning. In contrast to RV-specific PC occurrence in the intestines, neither general mLN hypertrophy, nor RV-specific CD8 T cell immunity are significantly impacted by components within immune breastmilk or the absence of protection to infection. Delaying onset of pup-intrinsic intestinal IgA secretion in the presence of active maternal protection may have profound benefits for the growing organism by conserving energy and/or allowing the early-life microbiome to colonize the gut.

Both breastmilk IgA and IgG were shown to contribute to the early life regulation of humoral immunity to the microbiota through mechanisms including immune exclusion and antibody-mediated immune regulation.^4,8,9^ The individual contributions of antibody isotypes is hard to disentangle in such a diverse system, especially given overlapping specificities within intestinal antibody repertoires and their profound effects on the microbiota in early life.^7,32,33^ Using murine RV infection enabled us to track the kinetics of neonatal immune onset in an antigen-specific manner, specifically probing for the individual effects of antigen-specific IgA and IgG provided by immune dams. As expected, protection from infection was mediated by IgA.^17^ This differs from maternal protection from *Citrobacter rodentium* infection, which has been shown to be mediated by breastmilk IgG.^28^ Since RV-specific IgA^+^ responses in the absence of maternal IgG were not delayed despite maintained protection from Rotavirus infection, this infection model allowed us to uncouple viral load from humoral immune response onset and instead suggest that breastmilk IgG fulfills an active and non-redundant role in shaping early-life humoral immunity.

Epitope masking represents a likely contributing mechanism for the delay in RV-specific B cell responses. In the adult system, antibody-mediated feedback shapes B cell selection and promotes epitope spreading, but does not completely block immune responsiveness to epitopes.29,34–36 Indeed, we found that PCs generated in the context of maternal protection showed overall lower clonal diversity, but similar levels of somatic hypermutation. This suggests that RV-derived antigens are retained in the pup in limited quantities and immune induction or clonal expansion towards the antigens can be initiated post weaning. While the delay in humoral onset is driven by IgG, the mechanism underlying decreased RV-specific clonal complexity in maternally protected offspring remains unclear. These two phenomena may or may not represent linked manifestations of maternal antibody mediated antigen limitation restricting the timing and breadth of GC founder recruitment while leaving affinity maturation intact. Regardless, the post-weaning established humoral immune memory was strong enough to shield maternally protected mice from reinfection during adulthood.

Seemingly contradicting our findings, a recent study using attenuated live Rotavirus as a model for vaccination showed that pre-existing maternal immunity completely blocked endogenous immune formation in the pup, rather than merely delaying the response.20 One difference could be original antigenic load, which may be higher in our case, using the WT, highly infectious EC_W_ Rotavirus strain. Another possible explanation could be that infection, even if blocked to levels undetectable by viral shedding quantification, alters the intestinal immune landscape, facilitating the integration of immune inducing signals. Along those lines, a recent study showed that breastmilk IgG from house dust mite (HDM)-immune dams, retrogradely taken in by pups via the neonatal Fc receptor expressed on the apical side of the intestinal wall, guided the development of allergic asthma to HDM only if the pups were exposed to HDM in the context of a pulmonary virus infection.31 Together, we propose that an immune activation threshold must be reached for the neonatal immune system to retain antigenic information for humoral immune response induction post weaning, and that this threshold is elevated by pre-existing maternal immunity through antigen-specific IgG.

Peri-weaning requirements for IgA response induction are poorly understood. The intestinal lamina propria of unperturbed pups harbors only few PCs, which increase dramatically in numbers post weaning, following the withdrawal from breastmilk. Given those findings, we wondered whether a mere lack of sufficient immune activation in protected pups or the mammary gland tissue of dams nursing infected pups^26^ could be the cause for the absence of local RV-specific IgA^+^ PC occurrence in the SILP prior to weaning. Importantly however, several of our observations argue against this notion: First, GC B cells specific to RV are equally diminished as those at effector sites in the context of maternal protection; second, RV-specific IgA^+^ PC induction in pups from RV-immune IgA-deficient dams remains delayed despite a lack of protection, and third, using a time-stamping approach labeling activated B cells and their progeny, we found that in contrast to those arising in the absence of maternal immunity, many of the RV-specific IgA^+^ PC clones found in maternally protected pups post-weaning did not originate from a classically 14 days post infection formed GC.

Early-life events significantly shape long-lasting homeostatic immunity and we have previously shown that Rotavirus infection in neonates increases the proportion of early-life derived adult IgA^+^ PCs in the SILP.^12^ Accordingly, we show here that neonatal RV infection enhances pre-weaning total IgA^+^ PC counts, which is blunted if the pups were maternally protected from infection. While this could partially be explained by a simultaneous lack of PCs reactive to other RV proteins than VP6, this is unlikely to account for the total effect given the strong immunodominance to VP6 in this model^37^, and rather points to an increase of total humoral reactivity in infected pups. This suggests that increased total IgA^+^ PC occurrence in unprotected infected pups pre-weaning is due to an infection-induced barrier breach by luminal antigens, as detected in the absence of total maternal antibody.^4^ Alternatively, changes to the intestinal wall might occur upon infection, which support the recruitment and/or retention of IgA^+^ PCs. Indeed, long-lived RV-specific IgA^+^ SILP PCs have recently been shown to depend on IFNγ-mediated upregulation of CXCR3 in adult mice, aiding their correct positioning at the effector site.^22^ The required type 1 immunity initiated in RV-infected mice might be provided by homeostatic mechanisms generally referred to as the “weaning reaction” in maternally protected pups, facilitating post-weaning humoral immune response initiation and long-lived IgA^+^ PC retention in the SILP.^11,22^

Oral immunization represents an attractive vaccination route, and breastmilk-mediated effects likely interfere with this immunization route in a unique manner. In contrast to previous studies interrogating the role of maternal protection on immune induction in the offspring using a systemic vaccination approach^38,39^, we here used an oral infection model. Across models and irrespective of the immunization route, pre-existing maternal immunity blocked pre-weaning humoral immune onset in an antigen-specific manner with surprising efficacy (^20,38,39^ and this study); however the long-term immune priming we describe here set our study apart from previous observations. Since the effects found in our study rely on breastmilk and we find substantial uptake of IgG1 through the intestinal wall, we propose that the positive effects conveyed by immune milk might depend on oral antigen administration and possibly reflect a general mechanism by which the growing infant, through breastmilk, receives not only passively protecting antibody, but also antigen-specific antibodies that help shape the long-term repertoire establishment. Based on our study and recent findings by others^9^, we propose that breastmilk IgG might function as an important rheostat to calibrate neonatal humoral immune responses to their necessary level, ensuring homeostasis and energy conservation alike. With that, our findings can explain how early-life derived triggers translate into long-lasting humoral immunity in the intestines, which is equally important for oral vaccine design as it is for the better understanding of how early life events guide long-lasting intestinal immune homeostasis.

## LIMITATIONS OF THE STUDY

We here use murine RV infection as a relevant model for neonatal enteric viral infection. One limitation of our study is that we cannot currently disentangle whether viral replication in maternally protected pups under the detection limit, active uptake of viral particles through the intestinal epithelium via breastmilk IgG, or both, are required for the described post-weaning onset of RV-specific humoral immunity in maternally protected pups.

Another limitation of the study is the lack of human data. Our study seemingly contrasts several previous reports suggesting that maternal protection limited immunity in newborns in response to vaccination. Maternal RV-specific IgA was shown to inversely correlate with pre-weaning seroconversion and RV-specific IgA levels in babies vaccinated with the oral attenuated live vaccine Rotarix^14–16^. In those studies, RV-specific IgA was routinely measured 4 weeks after the second vaccine dose, when babies were generally still breastfeeding. It would be highly interesting to test for fecal RV-specific IgA levels in vaccinated children at a later timepoint, as our study supports the hypothesis that protection from infection post weaning may not be impacted by maternal protection. Importantly however, isotype contributions to breastmilk antibody differ significantly between humans and mice, with IgG being much less prominent in human breastmilk. Whether our findings translate to the human system would be an important avenue of research guiding rational vaccine design.

## Material and Methods

### Key resources table

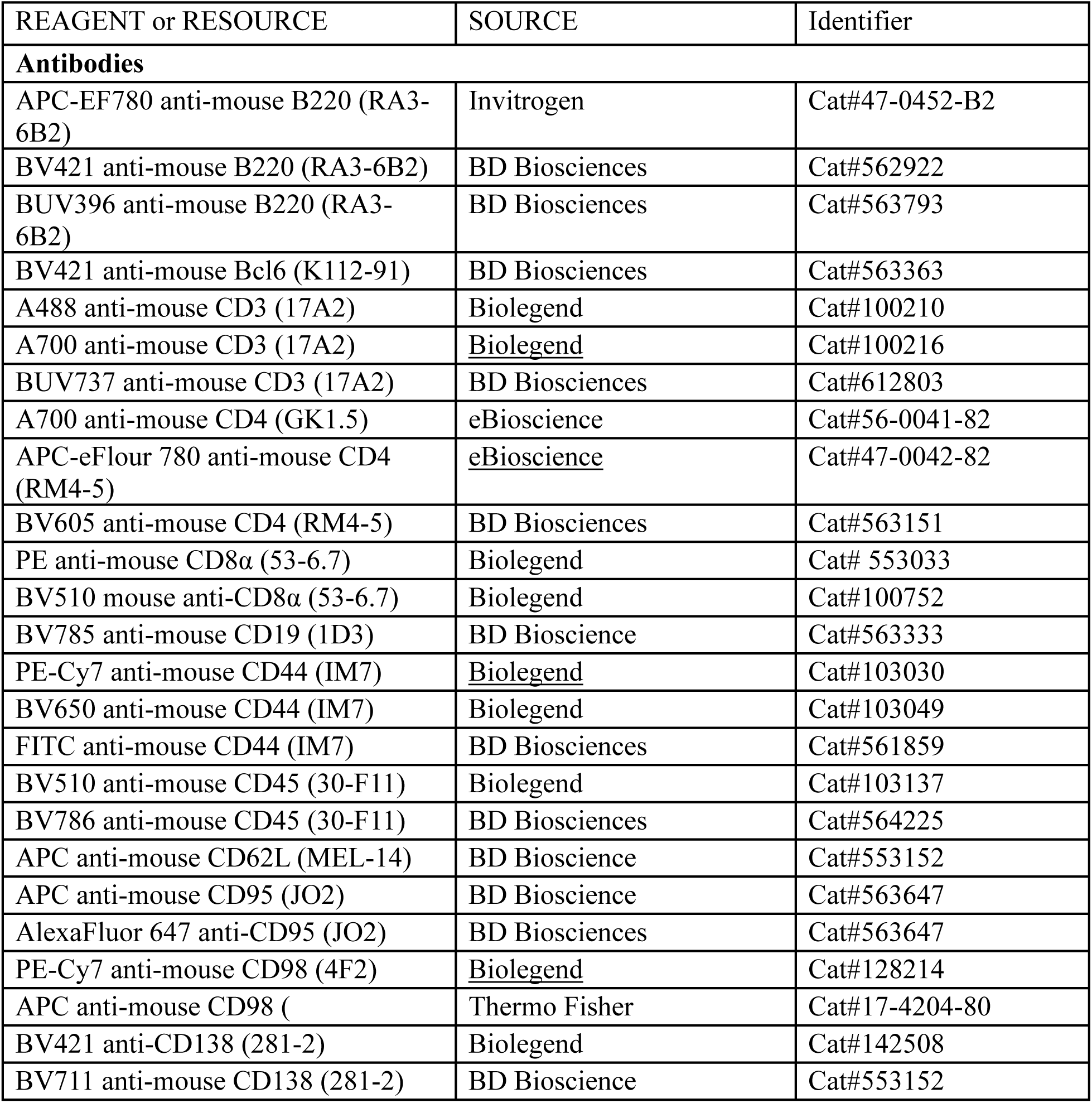
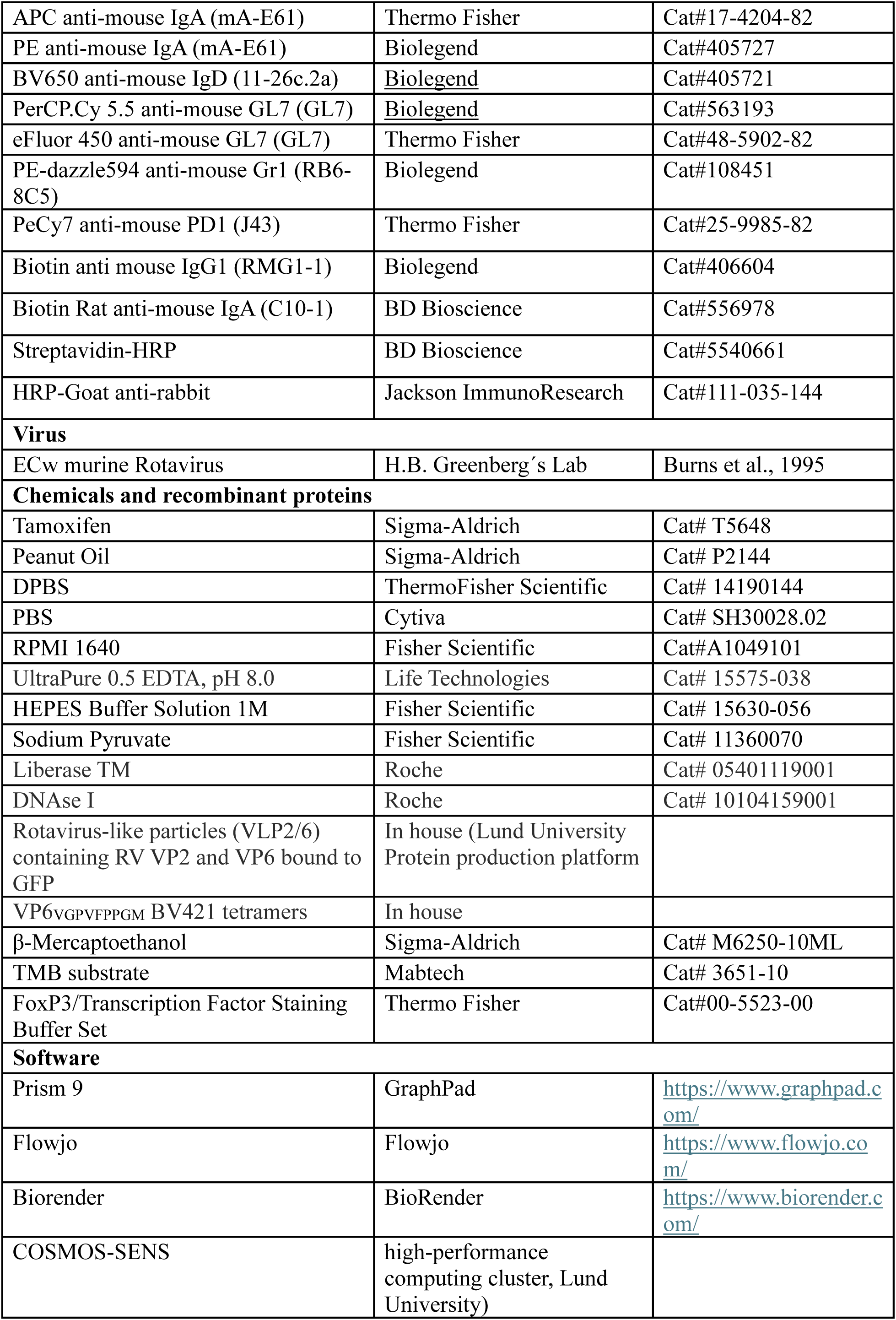
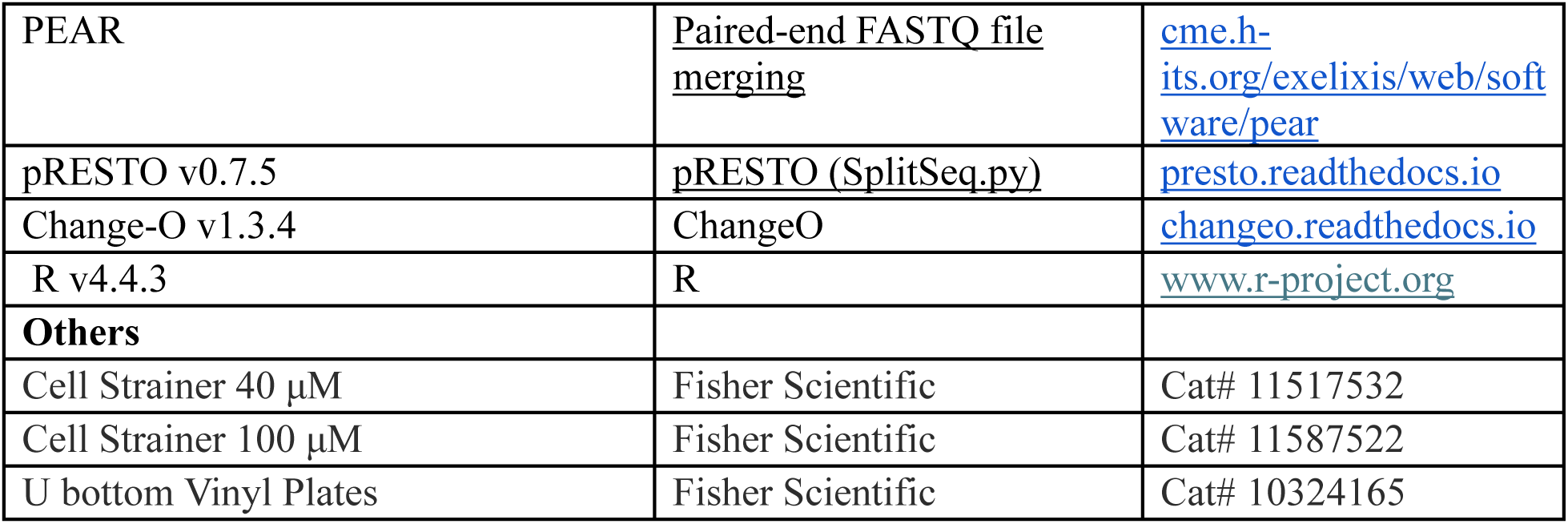

### Experimental model and subject details

#### Animals

All animals were housed under specific pathogen-free conditions at Lund, Sweden or the Spy Hill Campus at the University of Calgary, Canada. C57Bl/6NRj mice were purchased from Janvier labs or bred in house. µMT mice were purchased from The Jackson Laboratory and bred in house at the University of Calgary (JAX #002288, B6.129S2-Ighm^tm1Cgn^/J). IgA^27^ and IgG^28^ deficient mice were provided by Dr. Gabriel Nunez at the University of Michigan and maintained at the University of Calgary. *AID.Cre^ERT2^* mice (JAX 033897 *Aicda^tm1.1(cre/ERT2Crey)^*)^40^ were crossed to *R26^fl-STOP-Tom^* mice (JAX 007909 *Rosa26^LoxP-Stop-Lox-tdTomato^*)^41^ to generate time stamping mice as previously described.^12^ All animal breeding and procedures were performed in accordance with ethical permits approved by the Swedish Board of Agriculture (04525-17, 04765-22) or following the guidelines of the Canadian Council for Animal Care and approved by the University of Calgary Health Science Animal Care Committee (protocol #AC23-0172). Experimental groups consist of individual litters. Dams were littermates from heterozygous (dams) x homozygous (sires) maintenance colonies. Mice were maintained at an ambient temperature of ∼24 °C, humidity of ∼45%, and with 12-hour light-dark cycle.

### Method details

#### Rotavirus stock preparation

The virulent WT EC_w_ strain of RV stock was prepared by orally infecting 5 day old pups. Pups were euthanized two days post infection and intestines were harvested and FBS free M199 was added as 10% W/V. Tissues were homogenized with an electrically powered homogenizer and spun down at 3000 x *g*. Clear supernatants were used for the infections.

#### Rotavirus infection

Dams were infected with 5*10^4^ DD_50_ ECw virus in 100µl PBS by oral gavage at five to eight weeks of age and were set for breeding three weeks post infection. Experimental adult infection was performed at indicated timepoints. Neonatal mice were infected with 1*10^4^ DD_50_ ECw in 20µl volume by oral gavage on day 5 after birth. The mice were euthanized at the indicated time points for analysis.

#### Time stamping

For the induction of *AID.Cre^ERT2^* mediated time stamping, Tamoxifen (Sigma-Aldrich cat #T5648) was dissolved in peanut oil (Sigma-Aldrich, cat# P2144) at 30mg/ml and administered orally at a dose of 150mg/kg body weight on day 10 and 13 following rotavirus infection (corresponding to postnatal day 15 and 18, respectively).

#### Cell isolation

Small intestine and mesenteric lymph nodes were collected at indicated time points and single cell suspensions were prepared as previously described.^42^ Briefly, for SILP single cell suspensions, the intestinal content was flushed out with cold HBSS supplemented with 15mM Hepes followed by removal of Peyer’s Patches and fat. The intestine was longitudinally opened and cut into small pieces. Epithelial cells were removed by incubating the tissue for 15 min at 37°C with 2mM EDTA in HBSS supplemented with 10% FCS. This was followed by vigorous shaking and filtering using a nylon mesh. The EDTA washing was repeated once more for the younger mice (2wpi after d5 infection) and two more times for the older/adult mice, and the remaining tissue was digested in RPMI medium (Gibco cat#11875093) containing 10% FBS, 0.3 Wünsch/mL Liberase TM (Roche cat# 05401119001) and 30μg/ml DNase I (Roche cat#10104159001) on a magnetic stirrer for 15-20 mins at 37°C. The digested cell suspensions were filtered through a 100μm strainer (Fisher Scientific) and lymphocytes were enriched by density gradient centrifugation with 40%/70% Percoll (cytiva #17089101).

MLN cells suspensions were prepared by mashing the lymph nodes through a 70µm cell strainer in FACS buffer.

#### Flow cytometry

For the analysis of the cells by flow cytometry, staining was performed at a density of 1x10^7^ cells/100 μL after blocking for nonspecific binding with 10% rat serum (Sigma, cat#R9759) and/or rat α-mouse CD16/CD32 Fc block (BD, clone 2.4G2) for 20 min at 4°C in a FACS buffer. Dead cells were excluded using propidium iodide (Sigma–Aldrich, cat#537060), 7-Aminoactinomycin D (7AAD) ( Sigma–Aldrich, cat#A9400), or LIVE-DEAD Blue fixable dye and cell aggregates were excluded by identification on FSC-A versus FSC-H scatterplots. For the identification of virus specific B cells, the cells were stained for 50 min with virus like particles (VLP) containing RV-VP2 and VP6 bound to GFP prior to surface antibody staining. For the identification of RV-specific CD8 T cells, BV421-labeled tetramer containing the RV-VP6_VGPVFPPGM_ immunodominant peptide was preincubated at 37°C for 15 minutes, then the surface staining antibody cocktail was added and incubated for 30 minutes at 4°C in a FACS buffer as detailed previously.^43^ For identifying RV-specific CD4 T-cells, APC-labeled tetramer containing the RV-VP6_ATWYFNPVILRPNNV_ (from the NIH tetramer facility) was preincubated with the cell suspension for one hour at room temperature before surface and intracellular staining. Intranuclear targets were stained using a FoxP3/Transcription Factor Staining Buffer Set (Thermo Fisher) following the manufacturer’s protocol with the exception of the staining incubation which was performed overnight at 4°C. All FACS experiments were performed at the Lund University Immunology section, Lund Stem Cell Center FACS Core Facility, or the International Microbiome Centre (IMC) within the Snyder Institute at the University of Calgary on an LSRFortessa-X20, an LSRII (BD Biosciences), or a Cytek^®^ Aurora and analyzed using FlowJo software (BD).

#### ELISA

Fecal samples/colon content from infected and control mice were prepared by weighing and soaking in PBS, 1% BSA, 1 mM EDTA, soybean trypsin inhibitor (0.05 mg/ml), 2 mM PMSF phenylmethylsulfonyl fluoride) (Sigma) and 0.025% azide for 2 h at 100 mg/ml, homogenized and centrifuged at13000 rpm for 10 min. The supernatant was analyzed for the presence of RV antigen and RV/VP6-specific antibody as follows. Soft vinyl round bottom ELISA plates (Fisher scientific, Ref# 10324165) were coated with guinea pig anti-RV hyperimmune serum (kindly donated from Prof. Harry Greenberg, Stanford University) at 1:5000 in PBS and incubated at 37°C for 4hrs followed by blocking with 2% BSA in PBS for 2hrs at 37°C. Subsequently, fecal supernatants were diluted 1:20 in 0.5% BSA and added to plate and incubated at 4°C overnight. Rabbit anti-RV hyperimmune serum (from Prof. Harry Greenberǵs lab) was added (1:5000 in PBS) and incubated for 2hrs at 37°C. Detection was performed using HRP-Goat anti-rabbit antibody (Jackson Immuno research, Cat#111-035-144) for 1hr at 37°C. TMB substrate (Mabtech, Cat# 3651-10) was used to visualize the presence of the viral antigen.

RV-specific IgA and IgG1 detection in fecal supernatant, serum and stomach content (sample preparation for stomach content is the same as feces) was run similarly except that Rabbit anti-RV hyperimmune serum (kind gift from Prof. Harry Greenberg) was used as a capture antibody for Rhesus rotavirus (for total RV specific IgA/IgG1 detection) or purified VP6 (for VP6 specific IgA detection) that was incubated overnight at 4°C. After this, the plates were washed and samples were added at the desired dilutions (Feces supernatant 1:10, Serum 1:1600, stomach content supernatant 1:2). After overnight incubation at 4°C, IgA and IgG1 were detected using a biotinylated rat anti-mouse IgA (BD Bioscience, cat# 556978) or Biotin anti-mouse IgG1(BD Bioscience, cat# 406604), respectively. This was incubated for 2hrs at room temperature followed by washing and adding streptavidin-HRP (BD Bioscience, cat# 554088). After incubation for 1hr at room temperature, the plates were washed and TMB substrate (Mabtech, Cat# 3651-10) was used to visualize the presence of virus specific antibodies.

#### Bulk IgHA VDJ sequencing

VLP^+^IgA^+^ plasma cells (PCs) from the small intestinal lamina propria (SILP) were FACS-sorted from adult mice (n = 17; 5-8 weeks of age) that had been born to naïve or rotavirus (RV)-immune dams and infected neonatally with RV at day 5 after birth. Total RNA was extracted from sorted cells using the Quick RNA MicroPrep Kit (Zymo Research, R1050) according to the manufacturer’s instructions. Bulk IgHA VDJ sequencing (VDJ-seq) was performed as described previously.^44^ In brief, IgHA transcripts were captured via the constant region, and cDNA was generated by template-switch reverse transcription using SMARTScribe Reverse Transcriptase (Takara Bio, 639537). Template-switched cDNA libraries were PCR-amplified with Q5 High-Fidelity DNA Polymerase (New England Biolabs, M0491), purified using AMPure XP beads (Beckman Coulter, A63880), and quantified with a Qubit Fluorometer using the dsDNA HS Assay Kit (Fisher Scientific, 10606433). Up to 2.5 ng of amplified cDNA was indexed with the Illumina Nextera XT Index Kit v2 Set A (Illumina, 15052163). Final library quality was assessed on an Agilent 2100 Bioanalyzer using the High Sensitivity DNA Kit (Agilent, 5067-4626). Indexed libraries were pooled and sequenced on an Illumina MiSeq using the MiSeq Reagent Kit v3 (600 cycles; Illumina, MS-102-3003).

#### Bulk IgHA VDJ sequencing pre-processing

Paired-end FASTQ files were merged on an in-house high-performance computing cluster (COSMOS-SENS, Lund University) using PEAR v0.9.11. Merged FASTQ files were converted to FASTA format using the “SplitSeq.py” function from the pRESTO package v0.7.5. Sequences were submitted to the IMGT/HighV-QUEST web portal for germline assignment and alignment of IgH regions using default parameters. Aligned data were processed with the Change-O package v1.3.4 within the Immcantation framework. Databases were generated using “MakeDb.py”, unproductive rearrangements were removed with “ParseDb.py”, and germline reference sequences for VH, DH, and JH genes were reconstructed with “CreateGermlines.py” (all v1.3.4). Complementarity-determining region 3 (CDR3) sequence similarity and initial clonal clustering were computed with “DefineClones.py” v1.3.4.

#### Bulk IgHA VDJ sequencing analysis

All downstream analyses were performed in R v4.4.3 and GraphPad Prism v10. For clonal family assignment, sequences were clustered by identical VH and JH gene usage, junction length, and a Hamming distance threshold of 0.15 using “DefineClones.py” (Change-O v1.3.4); the Hamming distance cut-off was chosen after manual inspection of nearest-neighbor distance distributions generated with the shazam package v1.3.0. B cell clone size (fraction of reads per clone out of total reads per sample) was normalized to the number of sorted cells per sample. Mean somatic hypermutation (SHM) load per B cell clone in each sample and group (naïve vs immune dams) was calculated with the “observedMutations” function in shazam v1.3.0 across the entire IgH region and visualized using ggplot2 v4.0.2. Clonal diversity was quantified using Shannon’s diversity index implemented in the vegan package v2.7-2.

#### Statistical analysis and visualization

Statistical analysis and plotting was performed on Graphpad Prism 9 and 10. Fig. 2H was created on Python using the pandas module for data processing and the seaborn module for plotting. The specific statistical test methods used are noted in the corresponding figure legends. Figure describing experimental timelines were made using BioRender.

## ACKNOWLEDGEMENTS

We thank the Lahl, Yuan, and Agace labs for their input throughout the project, Elzbieta Eriksson (Lund University), Katrina Ellestad and Erisa Budo (University of Calgary) for support with mouse colony surveillance, Anna Fossum (The Lund Stem Cell Center FACS Core facility) and Karen Poon (University of Calgary) for FACS machine maintenance, and M. Greyson Christoforo for help with plotting data. We acknowledge Protein Production Sweden (PPS) for providing facilities and experimental support, and we would like to thank Wolfgang Knecht and Celeste Sele for assistance. PPS is funded by the Swedish Research Council as a national research infrastructure. Flow cytometry at the University of Calgary was done with the support from International Microbiome Centre (IMC), Snyder Institute, University of Calgary. The IMC is supported by the Cumming School of Medicine, University of Calgary, Western Economic Diversification (WED) and Alberta Economic Development and Trade (AEDT), Canada. The authors acknowledge support from the National Genomics Infrastructure in Stockholm funded by Science for Life Laboratory, the Knut and Alice Wallenberg Foundation and the Swedish Research Council, and NAISS for assistance with massively parallel sequencing and access to the UPPMAX computational infrastructure. The H-2K(b) VGPVFPPGM BV421 (anti-RV VP6 tetramer) was produced in house by Sara Suaréz Hernández in the laboratory of Sine Reker Hadrup.

Funding: K.G.M. was supported by Fysiografen and Stiftelsen Samariten. S.I.P. was supported by Eyes High, TRIANGLE, and CIHR postdoctoral funding programs. K.L. was supported by The Independent Research Fund Denmark (2034-00024A), The Swedish Research Council (2020-01977 and 2021-01385), the Canadian Institute of Health Research (GLB-192248 and 202409PJT-526453), the Natural Sciences and Engineering Research Council of Canada (NSERC; RGPIN-2025-06709), by a Snyder Institute Catalyst grant, and by start-up funding from the Department for Microbiology, Immunology and Infectious Diseases and the Snyder Institute for Chronic Diseases at the University of Calgary. The J.Y. laboratory was supported by the European Research Council (101125425), the Knut and Alice Wallenberg foundation (2024.0097, JY), the Swedish research council (2022-00617), and Svenska Sällskapet för Medicinsk Forskning.

## DATA AVAILABILITY

VDJ-seq data will be made available on GEO upon acceptance. All other data is available through the lead author upon reasonable request.

## AUTHOR CONTRIBUTIONS

K.G.M and K.L conceived and designed the study. K.G.M and S.I.P performed all the experiments and analysis unless otherwise indicated with the support of S.v.D, G.M, R.F, D.C.P, A.S.M, N.S, M.A and G.L. Fate-mapping experiments and VDJ-seq analysis were performed under the direction of J.Y. by K.G.M and G.M., J.Y and K.L were involved in critical discussions throughout. K.L wrote the manuscript with substantial writing contributions from J.Y and critical input from K.G.M and S.I.P. S.I.P generated the final figures. All authors provided feedback on the manuscript.

## FIGURE LEGENDS

**Supplementary Figure 1.**
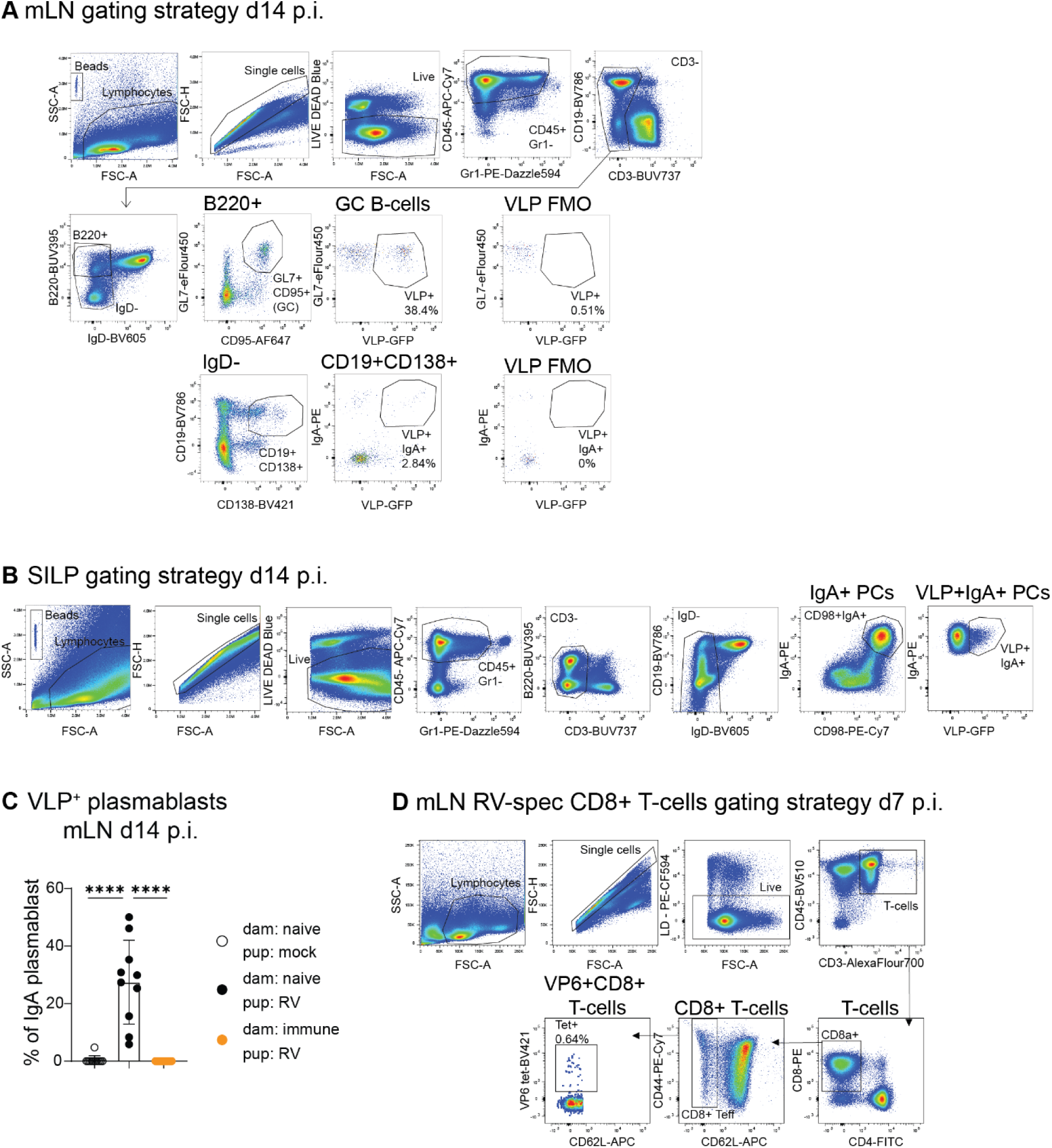
A) Representative mLN flow cytometry gating strategy to identify B cell subsets. B) Representative SILP flow cytometry gating strategy. C) Frequencies of VLP^+^ plasmablasts as a percentage of IgA^+^ plasmablasts in mLNs 2 weeks post neonatal day 5 RV infection of pups born to naïve (black) or immune (orange) dams. Naïve control (open circles). Bars indicate mean +/- SD. Statistical significance was determined using ordinary one-way ANOVA with Tukey’s multiple comparison test. ****p < 0.0001. D) Representative mLN flow cytometry to identify RV-specific CD8^+^ T-cells.

**Supplementary Figure 2.**
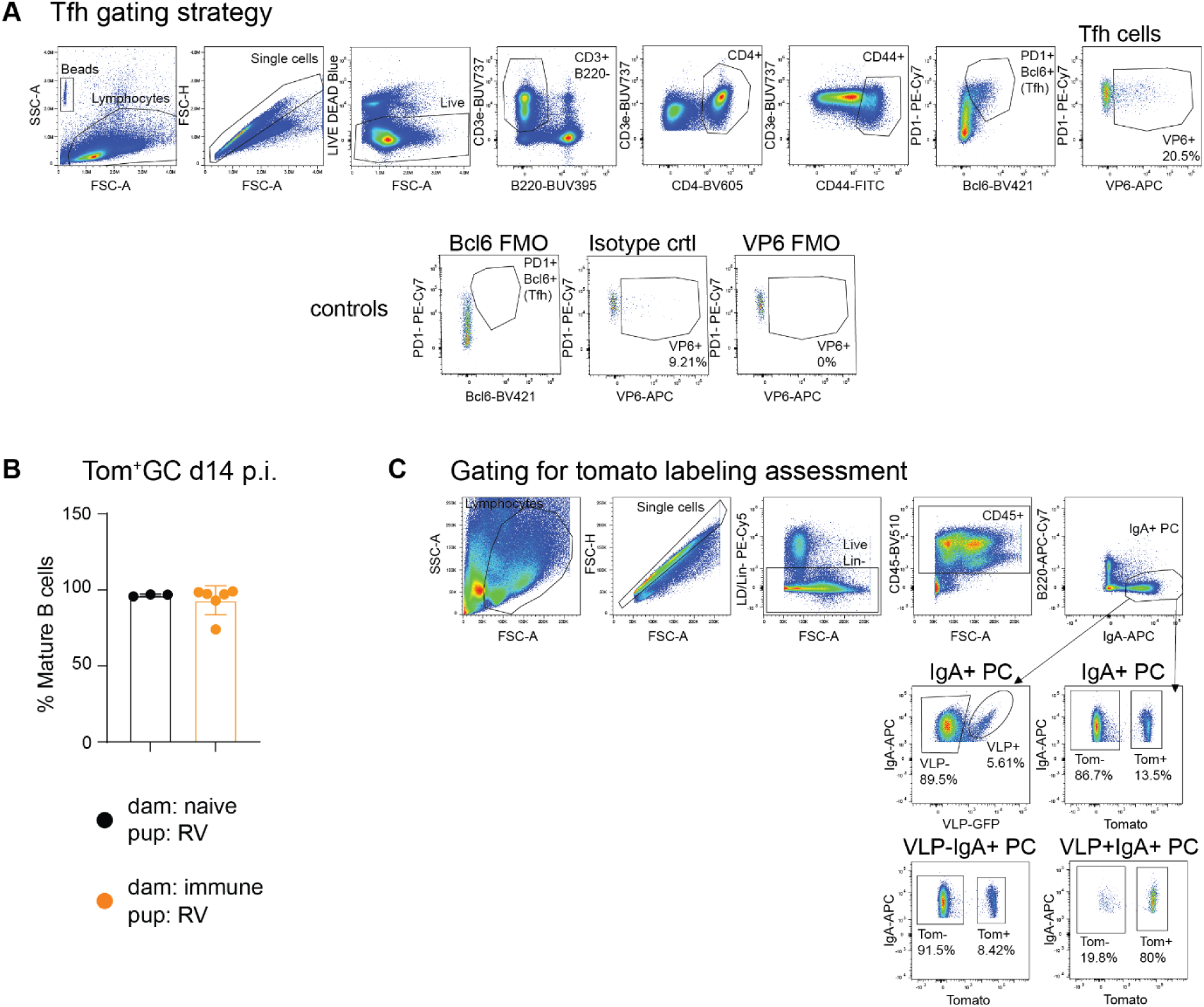
A) Representative mLN flow cytometry gating strategy to identify Tfh cells. B) Frequencies of Tomato^+^ GC B-cells as a percentage of mature B-cells in mLNs of AID.Cre^ERT2/WT^xTomato^LSL/WT^ pups born to naïve (black) or immune (orange) WT dams. Pups were orally infected with RV at neonatal day 5 and tamoxifen was administered at days 10 and 13 post infection. Baseline tomato labelling of GC B-cells was assessed one day after the last tamoxifen administration. The graph shows 1 experiment per group n = 3-6. Bars indicate mean +/- SD. Statistical significance was determined using a Two-tailed Mann-Whitney test. C) Representative SI and mLN gating strategy to identify Tomato^+^ cells.

**Supplementary Figure 3.**
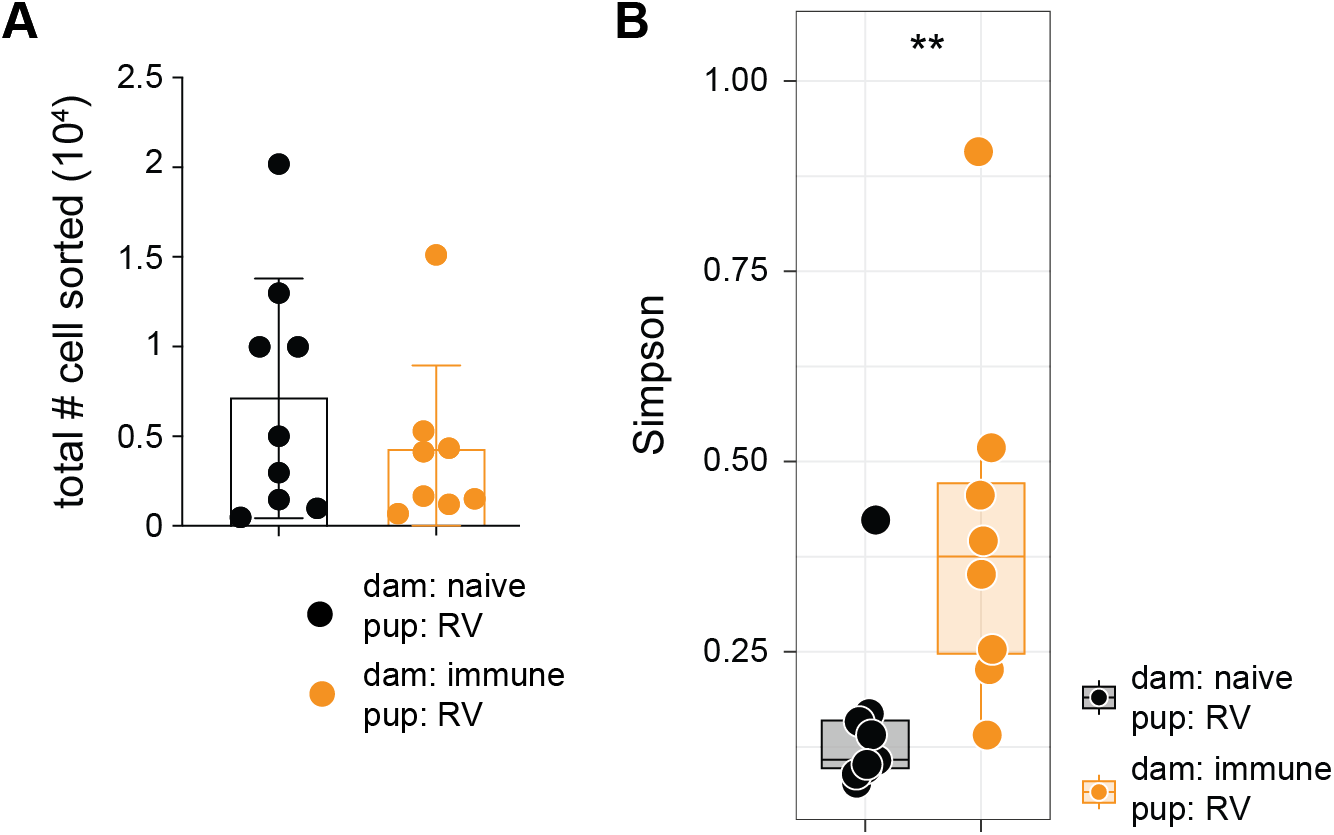
A) Quantification of total B cell numbers sorted for VDJ repertoire sequencing from adult offspring born to naïve (black, n= 9) or immune (orange, n= 8) dams following neonatal RV infection at day 5 post birth. Bars indicate mean +/- SEM. Statistical significance was determined using a two-tailed Wilcoxon rank-sum test, **p = 0.0078. B) Simpson Clonality Index calculated from clone frequencies within each sample (n = 9 adult offspring from naïve dams; n = 8 adult offspring from RV-immune dams), all infected neonatally with RV at day 5. Simpson Clonality is expressed as 1 − Simpson’s Diversity Index, where values approaching 1 indicate dominance of the repertoire by few clones. Each dot represents one biological replicate; box-and-whisker plots indicate median and interquartile range, with whiskers showing minimum and maximum values. Statistical significance was assessed using a two-tailed Wilcoxon rank-sum test **p < 0.01.

**Supplementary Figure 4.**
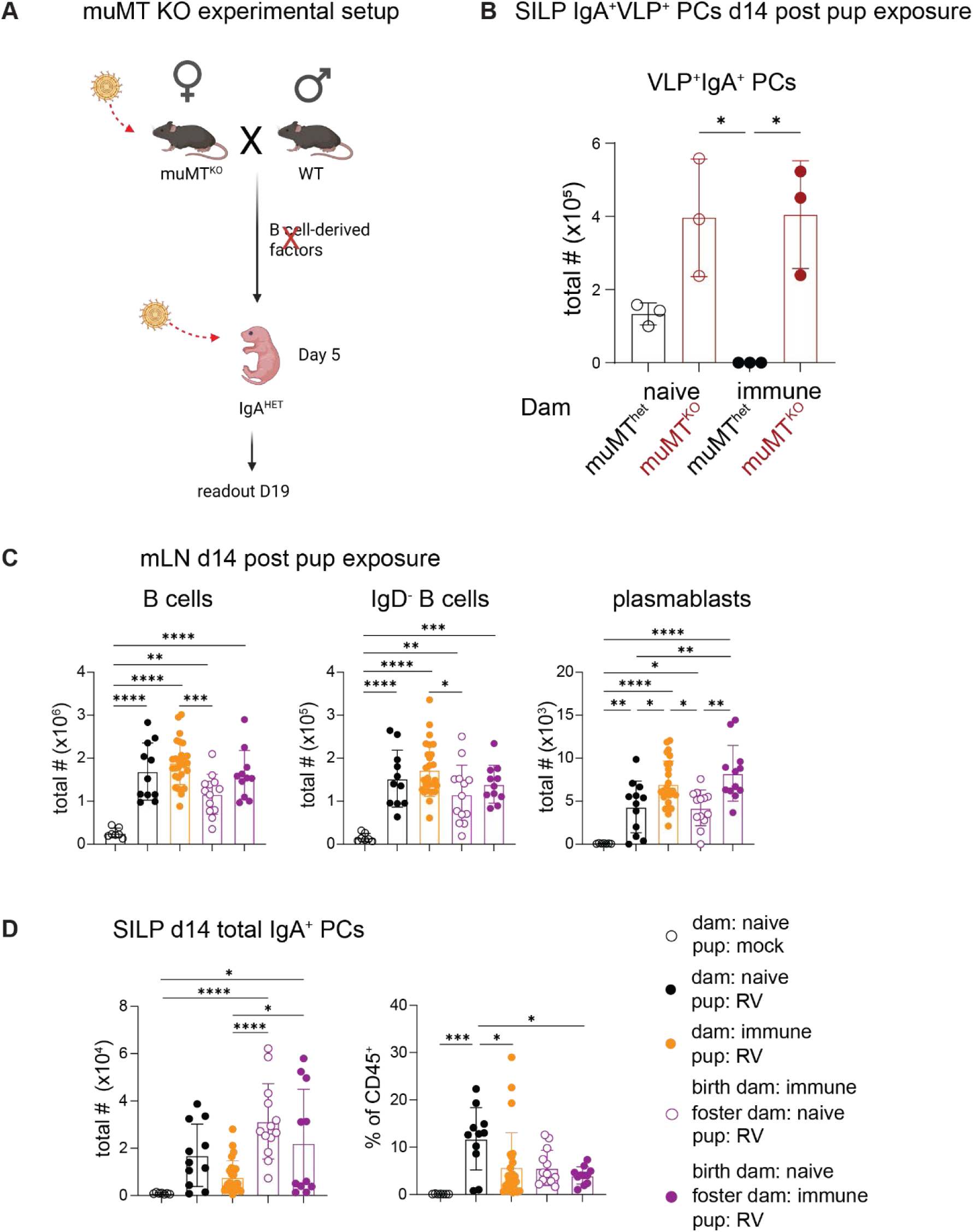
A) Experimental outline of experiments using µMT-deficient dams. B) Total numbers of VLP^+^IgA^+^ SILP PCs 2 weeks post neonatal day 5 oral RV infection of muMT HET/WT pups born to either naïve (open circles) or immune (closed circles) muMT HET (black) or muMT KO (red) dams. The graph shows one representative experiment of two with n=3/repeat. C) Mesenteric lymph node total B cell numbers (left, single cells, live CD45^+^CD19^+^), mature B cells (middle, single cells, live CD45^+^CD19^+^IgD^-^) and plasmablasts (right, single cells, live CD45^+^CD19^+^B220^-/low^IgD^-^CD138^+^) at 2 weeks post oral neonatal day 5 RV infection. Data is from mock (black open circles) or RV infected pups born to immune (orange) or mock (full black) infected dams. Cross-fostered litters (swapped within the first 24 hours post birth) were either born to immune and raised by naive (open purple circles) or born to naïve and raised by immune dams (full purple circles). The graphs show 2-5 independent experiments with n=2-8/repeat, except for uninfected control pups (1 experiment, n = 7 pups, black open circles). D) Total numbers (left) and frequencies (right) of SILP IgA^+^ PCs (single cells, live CD45^+^CD19^low/-^IgA^+^) analyzed two weeks post neonatal day 5 oral RV infection. Mice were the same as in C. Bars indicate mean +/- SD and statistical significance was determined using ordinary one-way ANOVA with Tukey’s multiple correction test. *p < 0.05, **p < 0.01, ***p < 0.001, ****p < 0.0001.

**Supplementary Figure 5.**
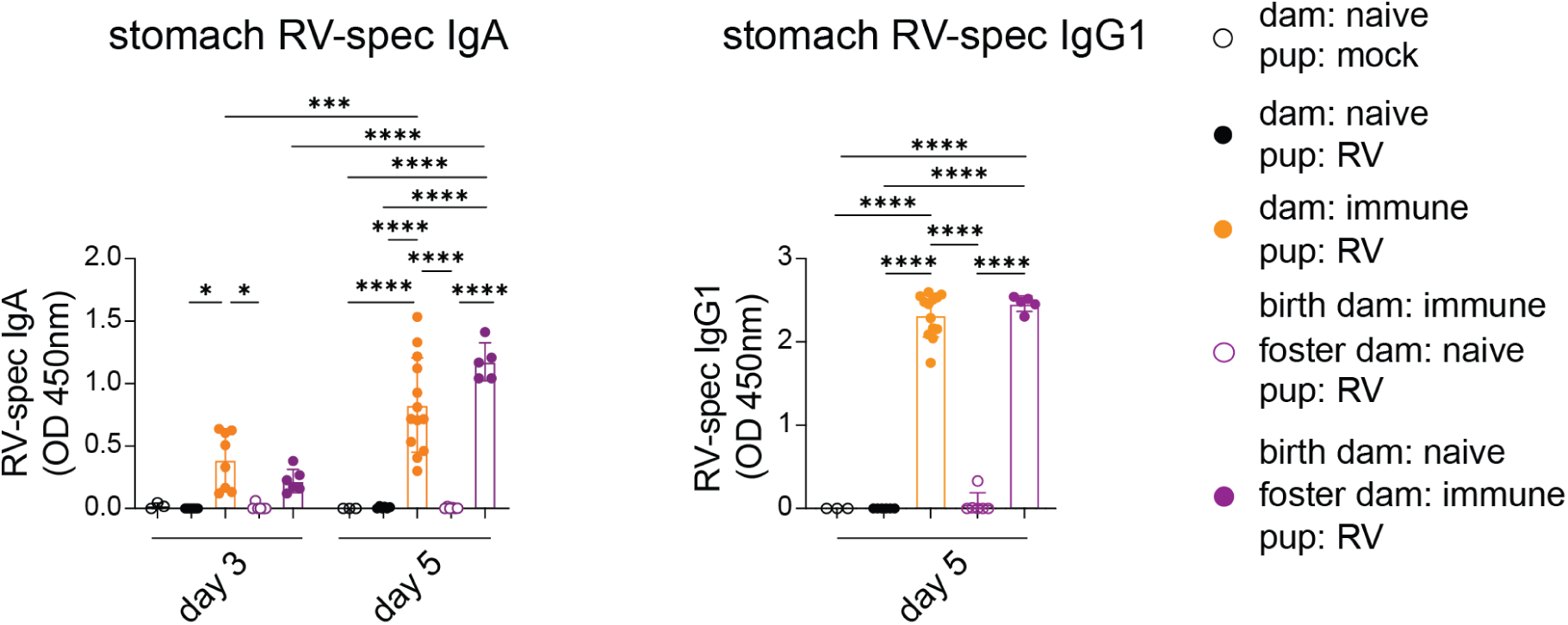
RV-specific IgA (left) and IgG1 (right) in the stomach content of mock (black open circles) or RV infected pups born to immune (orange) or mock (full black) dams at 3 and 5 days post infection for RV-specific IgA and at 5 days post infection for RV-IgG1. Cross fostered litters (swapped within the first 24 hours post birth) were either born to immune and raised by naive (open purple circles) or born to naïve and raised by immune (full purple circles) dams. The graph shows 2 independent experiments with (n= 2-8) except for naïve groups (1 experiment with n=3). Bars indicate mean +/- SD and statistical significance was determined using ordinary one-way ANOVA with Tukey’s multiple correction test. *p < 0.05, **p < 0.01, ***p < 0.001, ****p < 0.0001.

