## Supplementary for "Breastmilk antibody isotypes differentially protect against neonatal rotavirus infection and modulate the timing and clonal architecture of offspring B cell responses"

**Supplementary Figure 5**

RV-specific IgA (left) and IgG1 (right) in the stomach content of mock (black open circles) or RV infected pups born to immune (orange) or mock (full black) dams at 3 and 5 days post infection for RV-specific IgA and at 5 days post infection for RV-IgG1. Cross fostered litters (swapped within the first 24 hours post birth) were either born to immune and raised by naive (open purple circles) or born to naïve and raised by immune (full purple circles) dams. The graph shows 2 independent experiments with (n= 2-8) except for naïve groups (1 experiment with n=3). Bars indicate mean +/- SD and statistical significance was determined using ordinary one-way ANOVA with Tukey's multiple correction test. *$p < 0.05$, **$p < 0.01$, ***$p < 0.001$, ****$p < 0.0001$.

Supplementary Figure 1

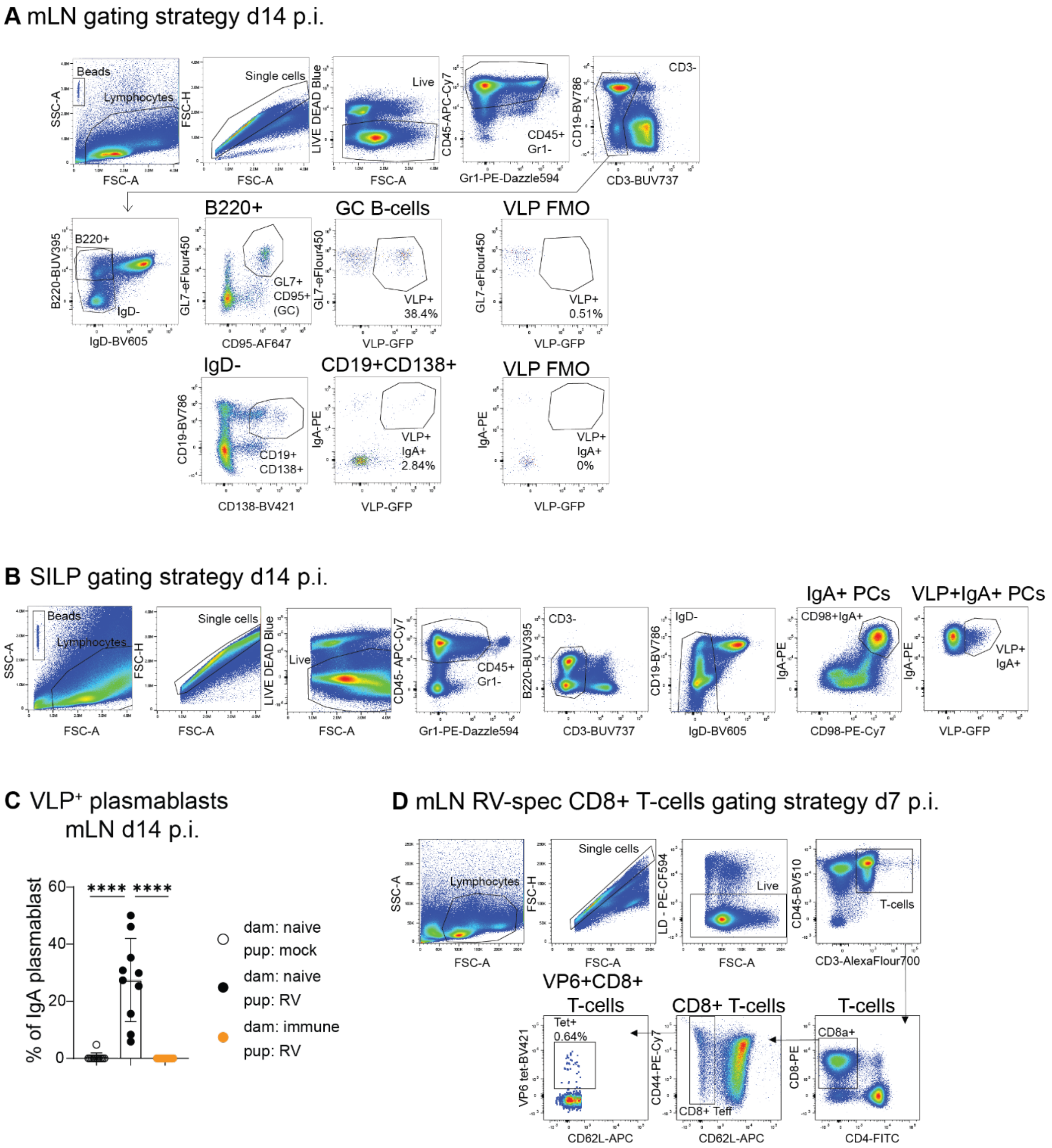

Supplementary Figure 2

**A** Tfh gating strategy

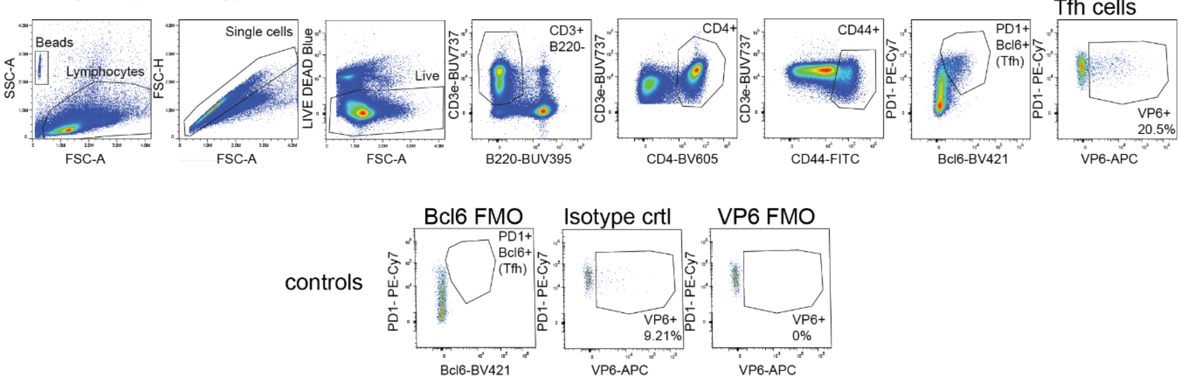


**B** Tom⁺GC d14 p.i.

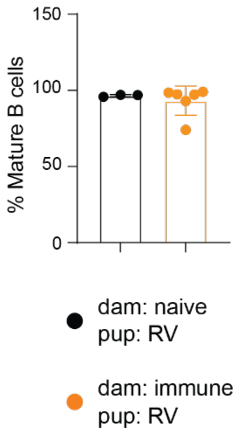


**C** Gating for tomato labeling assessment

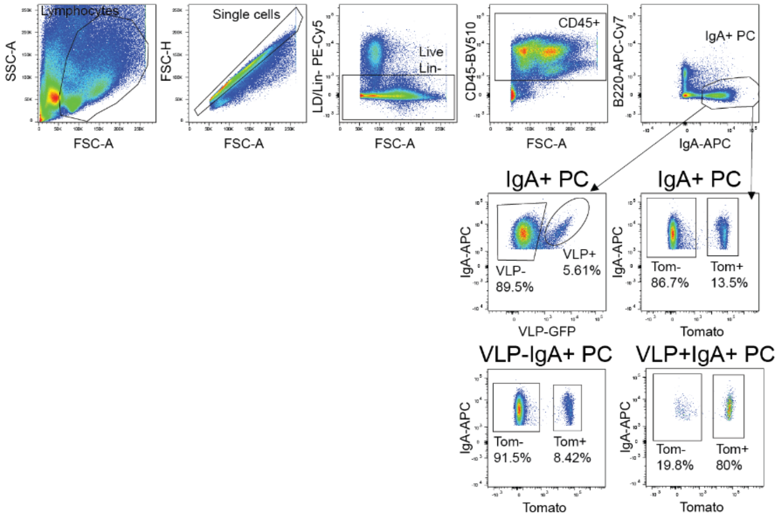

Supplementary Figure 3

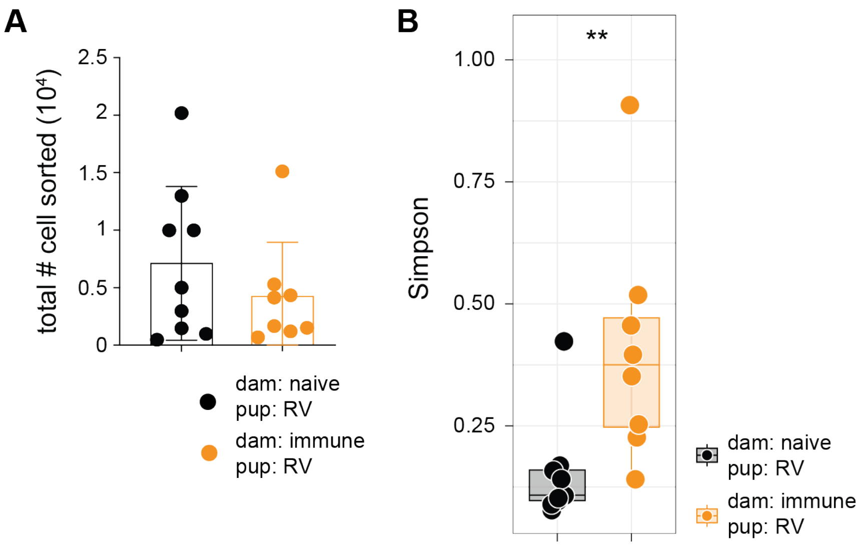

Supplementary Figure 4

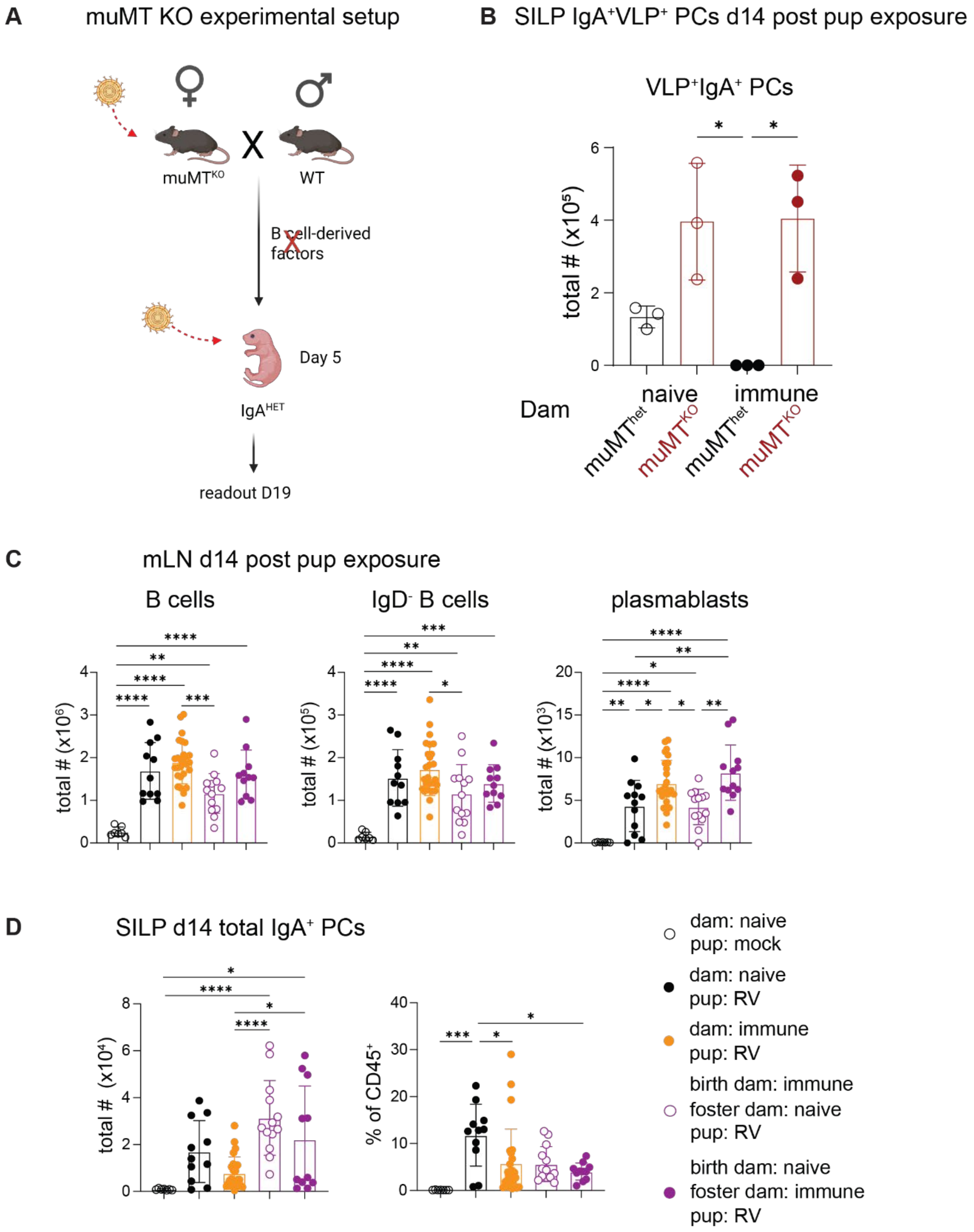

Supplementary figure 5

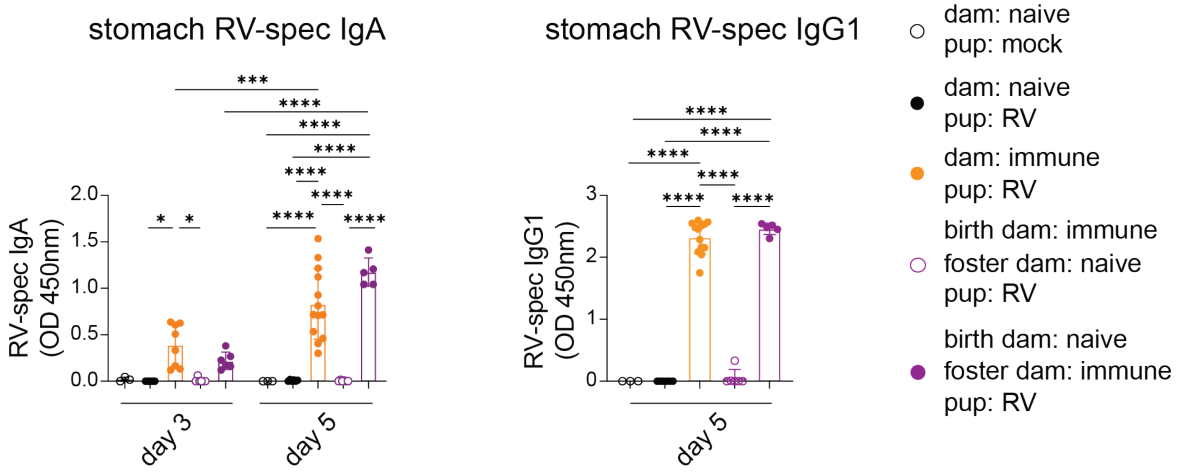
